# Dual CD40/CD28 co-stimulation reprograms the solid-tumor microenvironment to unleash CAR-T and endogenous anti-tumor immunity

**DOI:** 10.64898/2026.11.08.762395

**Authors:** Zhenguo Zi, Cong Han, Fei Yang, Jie Wang, Gengzheng Zhu, Xiaojun Liu, Yangbing Zhao

## Abstract

Chimeric antigen receptor (CAR) T-cell therapy has shown limited efficacy against solid tumors, owing to poor T-cell persistence, an immunosuppressive tumor microenvironment (TME) and cancer immune evasion. Here we describe LACO-Stim (lymphocytes-APC co-stimulator), a chimeric co-stimulatory molecule composed of an agonistic anti-CD40 single-chain variable fragment (scFv) and the CD28 intracellular signaling domain, such that a single molecule activates CD40-expressing antigen-presenting cells while delivering CD28 co-stimulation to the T cell that bears it. Co-expressed with a 4-1BB-based CAR (LACO-CAR-T), it enhanced anti-tumor activity in vitro and achieved durable tumor regression rather than transient control in xenograft and immunocompetent models. In immunocompetent hosts, LACO-CAR-T activated dendritic cells, reprogrammed tumor macrophages toward M1-like, antigen-presenting states, and induced epitope spreading with endogenous CD8^+^ T-cell responses against tumor neoantigens. Single-cell transcriptomics and RNA-velocity analysis revealed coordinated immune remodeling: T cells shifted toward a more stem-like, less terminally exhausted state, regulatory T cells declined in frequency and suppressive function, and myeloid cells acquired antigen-presentation programs while excessive inflammatory and suppressive signaling was restrained. LACO-Stim further synergized with anti-PD-1 blockade to control tumor growth. LACO-Stim thus enables CAR-T cells to coordinate intrinsic T-cell reinforcement with extrinsic immune reprogramming, converting an immunosuppressive TME into a de-inflamed (reduced inflammatory signaling), de-suppressed rather than merely armed niche that supports endogenous anti-tumor immunity.

## Introduction

Chimeric antigen receptor (CAR) T-cell therapy has transformed the treatment of hematological malignancies, yet its efficacy against solid tumors remains limited by three principal barriers: inadequate T-cell infiltration and persistence in the absence of sustained co-stimulatory signals, a profoundly immunosuppressive tumor microenvironment (TME) that restrains both engineered and endogenous anti-tumor immunity, and immune escape driven by tumor heterogeneity [1, 2]. Strategies developed to overcome these barriers have focused either on improving the intrinsic functionality of CAR-T cells, through optimized co-stimulatory domains, armored cytokine secretion or resistance to checkpoint signals, or on modulating the TME through combination therapies [2]. Few approaches, however, address both cell-intrinsic and cell-extrinsic mechanisms within a single molecular design.

CD40, a member of the tumor necrosis factor receptor superfamily, is a central regulator of antigen-presenting cell (APC) function [3–5]. Engagement of CD40 on dendritic cells (DCs) and macrophages promotes APC maturation, upregulates the co-stimulatory molecules CD80 and CD86 and major histocompatibility complex (MHC) molecules, and triggers secretion of pro-inflammatory cytokines including interleukin-12 (IL-12), collectively initiating robust adaptive anti-tumor responses [6]. Agonistic CD40 antibodies have shown anti-tumor activity in preclinical models and early-phase clinical trials, including pancreatic carcinoma [7]. CD40 signaling sustains this IL-12 production and promotes T helper 1 (TH1) and cytotoxic T lymphocyte (CTL) differentiation [8, 9], and CD40 engagement on APCs facilitates cross-presentation of tumor-associated antigens on MHC class I molecules to CD8^+^ T cells, driving epitope spreading beyond the engineered specificity [10–12]. In macrophages, CD40 engagement promotes fatty acid oxidation and glutamine metabolism, which support the epigenetic reprogramming required for the transition from an M2-like, immunosuppressive to an M1-like, immunostimulatory state, thereby restoring antigen presentation, recruiting and activating CD8^+^ T and NK cells, and converting the TME into a tumor-rejecting niche [5, 7, 13].

CD28 provides the second signal that T cells require to convert antigen recognition into effector function. In the TME, CD28 engagement by CD80/CD86 on APCs, including reprogrammed tumor-associated macrophages (TAMs), amplifies TCR-proximal signaling through PI3K–AKT and Ras–MAPK, lowering the activation threshold and imposing transcriptional and epigenetic programs that drive IL-2 production, proliferation, survival and effector differentiation while preventing anergy and exhaustion. Because CD28 is also the preferred dephosphorylation target of PD-1– recruited SHP2, restoring CD28 co-stimulation relieves PD-L1/PD-1-mediated suppression, enabling endogenous and CAR-T cells to regain cytotoxic and infiltration capacity in an otherwise immunosuppressive TME [14–16].

Whether deliberately co-delivering both CD40 and CD28 signals in the TME can reprogram the solid-tumor microenvironment to unleash CAR-T and endogenous anti-tumor immunity remains untested. We therefore designed LACO-Stim (lymphocytes-APC co-stimulator), a single chimeric molecule that couples a functionally validated agonistic anti-CD40 scFv to the CD28 intracellular signaling domain: unlike membrane-bound CD40 ligand [17], the anti-CD40 arm does not require ligand-like trimerization to engage CD40, while the CD28 arm delivers co-stimulation in cis to the co-expressed CAR-T cell, enabling it to resist TME-mediated suppression. Here we show that LACO-CAR-T confers durable tumor control in xenograft and immunocompetent models, licenses APCs within the tumor, elicits epitope spreading and endogenous CD8^+^ T-cell responses, and synergizes with PD-1 blockade.

## Results

### LACO-Stim activates dendritic cells and polarizes macrophages in vitro and potentiates CAR-T-cell function in immunodeficient mice

To determine whether LACO-Stim-expressing CAR-T cells activate CD40-expressing APCs, we differentiated monocytes from healthy donors into dendritic cells (DCs) or macrophages (Mφs). A lentiviral vector encoding LACO-Stim, comprising an agonistic anti-human CD40 scFv fused to the human CD8α hinge/transmembrane domain and the human CD28 intracellular domain, was co-expressed with an IL13Rα2 CAR containing 4–1BB and CD3ζ intracellular domains (Fig. 1a). Co-incubation of DCs with LACO-CAR-T cells for 24 h upregulated CD80, CD86, CD83 and HLA-DR (Fig. 1b) and increased secretion of IL-12, GM-CSF and IL-1β (Fig. 1c) relative to DCs co-incubated with conventional CAR-T cells, indicating that the CD40 signal delivered by LACO-Stim drives DC maturation and activation. Similarly, Mφs co-cultured with LACO-CAR-T cells showed increased CD80, CD86 and HLA-DR, reduced expression of the M2-associated markers CD206 and CD163 (Fig. 1d) and increased secretion of IL-6, GM-CSF and IL-1β (Fig. 1e), consistent with M1-like polarization. LACO-CAR-T cells also significantly enhanced Mφ phagocytic capacity when co-cultured with targeted tumor cells compared with CAR-T cells, whereas non-transduced (NTD) T cells did not affect macrophage phagocytosis (Fig. 1f, g). These results establish that LACO-Stim activates CD40-expressing myeloid cells through its agonistic anti-CD40 scFv domain, promoting DC maturation and reprogramming Mφs toward a pro-inflammatory phenotype.

**Fig. 1.**
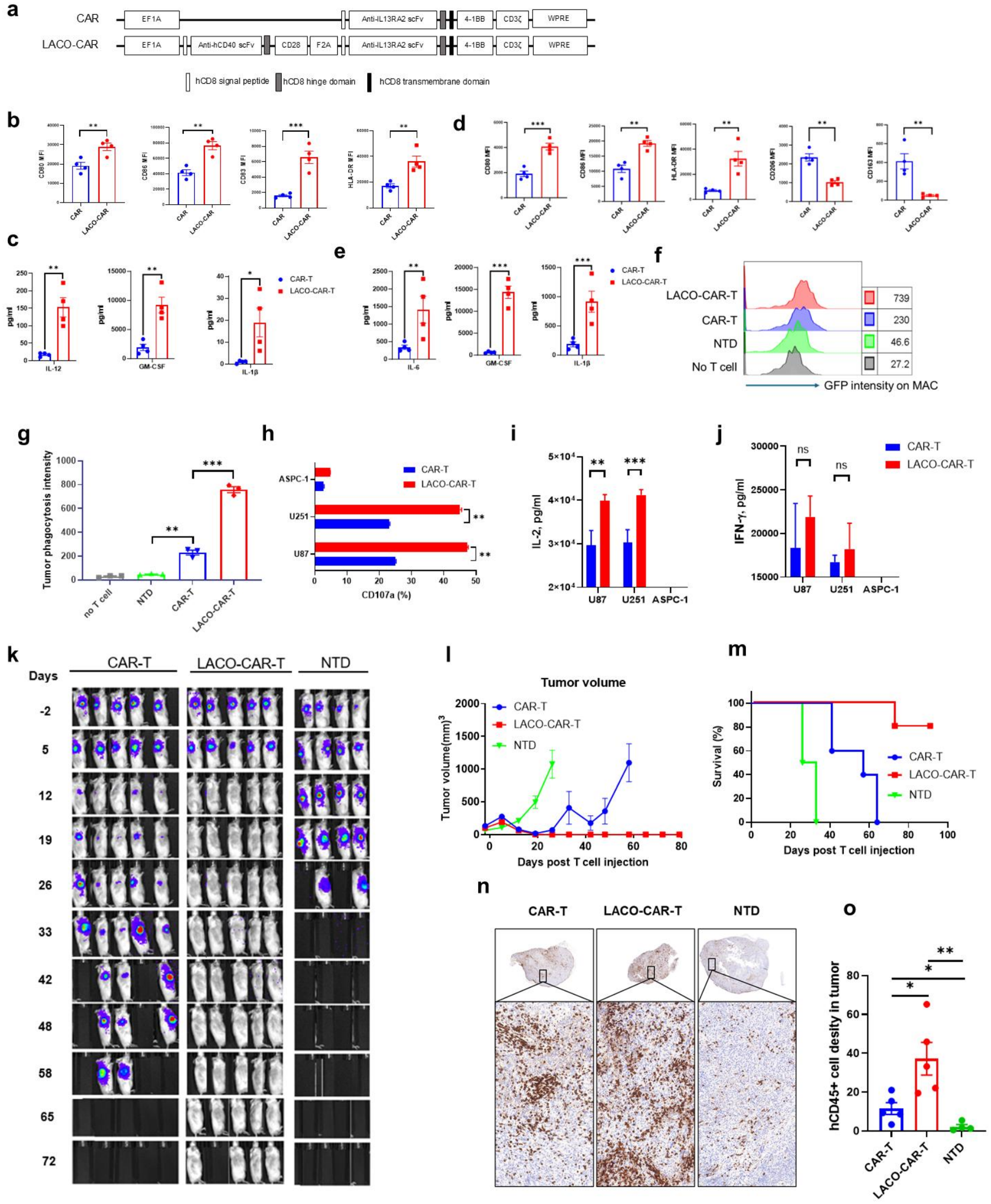
LACO-Stim activates dendritic cells and polarizes macrophages toward an M1-like phenotype and enhances CAR-T-cell function in vitro and in vivo. a, Schematic of the LACO-CAR and CAR constructs. b, Flow-cytometry analysis of CD80, CD86, CD83 and HLA-DR expression on monocyte-derived dendritic cells (DCs) after co-culture with CAR-T or LACO-CAR-T cells for 24 h, with statistical analysis of four individual donors (two-way ANOVA; *P < 0.05, **P < 0.01). c, Cytokine secretion (IL-12, GM-CSF and IL-1β) in supernatants after co-culture of DCs with CAR-T or LACO-CAR-T cells for 24 h. d, Flow-cytometry analysis of CD80, CD86, HLA-DR, CD206 and CD163 expression on monocyte-derived macrophages (Mφs) after co-culture with CAR-T or LACO-CAR-T cells for 24 h, with statistical analysis of four individual donors (two-way ANOVA; *P < 0.05, **P < 0.01, ***P < 0.001). e, Cytokine secretion (IL-6, GM-CSF and IL-1β) in supernatants after co-culture of Mφs with CAR-T or LACO-CAR-T cells for 24 h. f, Representative flow-cytometry analysis of GFP intensity in Mφs after co-culture with U87-CBG cells in the presence of CAR-T, LACO-CAR-T or NTD cells for 18 h. A total of 1 × 10⁵ Mφs were co-cultured with 2.5 × 10⁴ T cells and 1 × 10⁵ U87-CBG cells in 96-well flat-bottom plates; green fluorescence signals (originating from tumor cells) were detected in Mφs by FACS with gating on the 7-AAD⁻CD45^+^CD3⁻ population. g, Statistical analysis of green fluorescence intensity in Mφs in each group. Data are mean ± s.e.m. of three independent replicates; two-way ANOVA. **P < 0.01, ***P < 0.001. h, CD107a surface expression detected by flow cytometry at 4 h after co-culture of 1 × 10⁵ CAR-T or LACO-CAR-T cells with 1 × 10⁵ tumor cells in the presence of GolgiStop (BD Biosciences). Data are mean ± s.e.m. of three independent replicates; two-way ANOVA. **P < 0.01. i, j, ELISA quantification of IL-2 (i) and IFN-γ (j) secretion at 24 h after co-culture of 1 × 10⁵ CAR-T or LACO-CAR-T cells with 1 × 10⁵ tumor cells; two-way ANOVA. **P < 0.01, ***P < 0.001; ns, not significant (P > 0.05). k, Bioluminescence imaging of anti-tumor effects of CAR-T, LACO-CAR-T or NTD T cells on U87-CBG tumor-bearing mice using the IVIS system every 7 days until day 72 after treatment. l, Tumor volume measured by calipers until day 80 after treatment. m, Survival curves per group until day 90 after treatment. n, Representative CD45 immunohistochemistry of tumors from CAR-T, LACO-CAR-T and NTD groups at day 12 after treatment (n = 4 mice). o, Statistical analysis of tumor CD45 intensity per group (two-way ANOVA; *P < 0.05, **P < 0.01).

Two IL13Rα2-positive glioma lines (U87 and U251) served as targets, with ASPC-1 as a negative control. CD40 was detected on U87 but not on U251 or ASPC-1 cells (Extended Data Fig. 1a). Both IL13Rα2 CAR-T and LACO-CAR-T cells lysed U87 and U251 cells but not ASPC-1 cells (Extended Data Fig. 1b–d). In a CD107a degranulation assay, LACO-CAR-T cells upregulated CD107a to a greater extent than CAR-T cells upon co-culture with U87 and U251 (Fig. 1h). LACO-CAR-T cells also secreted more IL-2, but not IFN-γ, than CAR-T cells (Fig. 1i, j). In a U87-CBG xenograft NSG mouse model, mice treated with LACO-CAR-T cells maintained complete tumor regression for more than 80 days without evidence of relapse, with significantly enhanced survival (Fig. 1k–m). In a separate experiment, tumors collected 12 days after treatment were subjected to immunohistochemical staining for human CD45. Tumors in the LACO-CAR-T treatment group showed markedly increased T-cell infiltration relative to those in the CAR-T treatment group (Fig. 1n, o). Collectively, these data indicate that LACO-Stim-modified CAR-T cells exhibit superior anti-tumor potency against solid tumors in the NSG immunodeficient model.

### LACO-Stim recruits endogenous T cells and achieves durable tumor control in immunocompetent hosts

Because immunodeficient tumor models cannot fully recapitulate the effects of LACO-Stim CAR-T cells, we generated murine versions of the CAR and LACO-Stim by replacing the CD28 sequence of LACO-Stim and the CD8α, CD28, 4–1BB and CD3ζ sequences of the CAR with murine counterparts. MC38 cell lines expressing human IL13Rα2 or MSLN engrafted poorly in C57BL/6 mice; we therefore used human CD19 (hCD19) as a surrogate antigen and generated MC38-hCD19 tumor lines and an anti-hCD19 (FMC63) CAR (Fig. 2a). Because the anti-hCD40 scFv does not cross-react with murine CD40, we used human CD40 humanized transgenic C57BL/6 mice (hCD40/CD45.2), in which normal lymphoid development was confirmed (Extended Data Fig. 2a). hCD40/CD45.2 mice bearing established MC38-hCD19 tumors received sublethal lymphodepleting irradiation (4.75 Gy) on day 13 after tumor inoculation, followed by intravenous transfer of CD45.1 transgenic murine T cells retrovirally transduced with the CD19 CAR or LACO-CAR (Extended Data Fig. 2b) on day 0 (day 14 after tumor inoculation) (Fig. 2b). LACO-CAR-T cells induced complete tumor regression with durable cures in most treated mice (5 of 7) (Fig. 2c) and significantly prolonged survival (Fig. 2e), whereas CAR-T cells alone transiently controlled tumor growth but failed to prevent recurrence on day 35 after T cell treatment (Fig. 2d). No body-weight loss was observed, indicating favorable tolerability (Extended Data Fig. 2c). In a separate experiment, tumors were collected on day 30 for immunophenotypic analysis (Extended Data Fig. 2d). LACO-CAR-T treatment substantially increased tumor-infiltrating T cells (Fig. 2f). Moreover, PD-1 expression on both exogenous (CD45.1^+^) and endogenous (CD45.2^+^) CD8^+^ T cells was significantly upregulated (Fig. 2g, h), suggesting that LACO-Stim induced activation of both exogenous and endogenous CD8^+^ T cells, and that tumor control was mediated jointly by infused and endogenous T-cell compartments. Although DC and Mφ phenotypes in blood and spleen were similar between the two groups (Extended Data Fig. 2e), CD86 and MHC-II expression were significantly increased on tumoral DCs of the LACO-CAR-T group (Fig. 2i, j), and intracellular IL-12 expression in tumoral DCs was significantly increased (Fig. 2k), suggesting that LACO-Stim enhanced the inflammatory state of tumoral DCs and promoted the recruitment of effective anti-tumor T cells into the tumor.

**Fig. 2.**
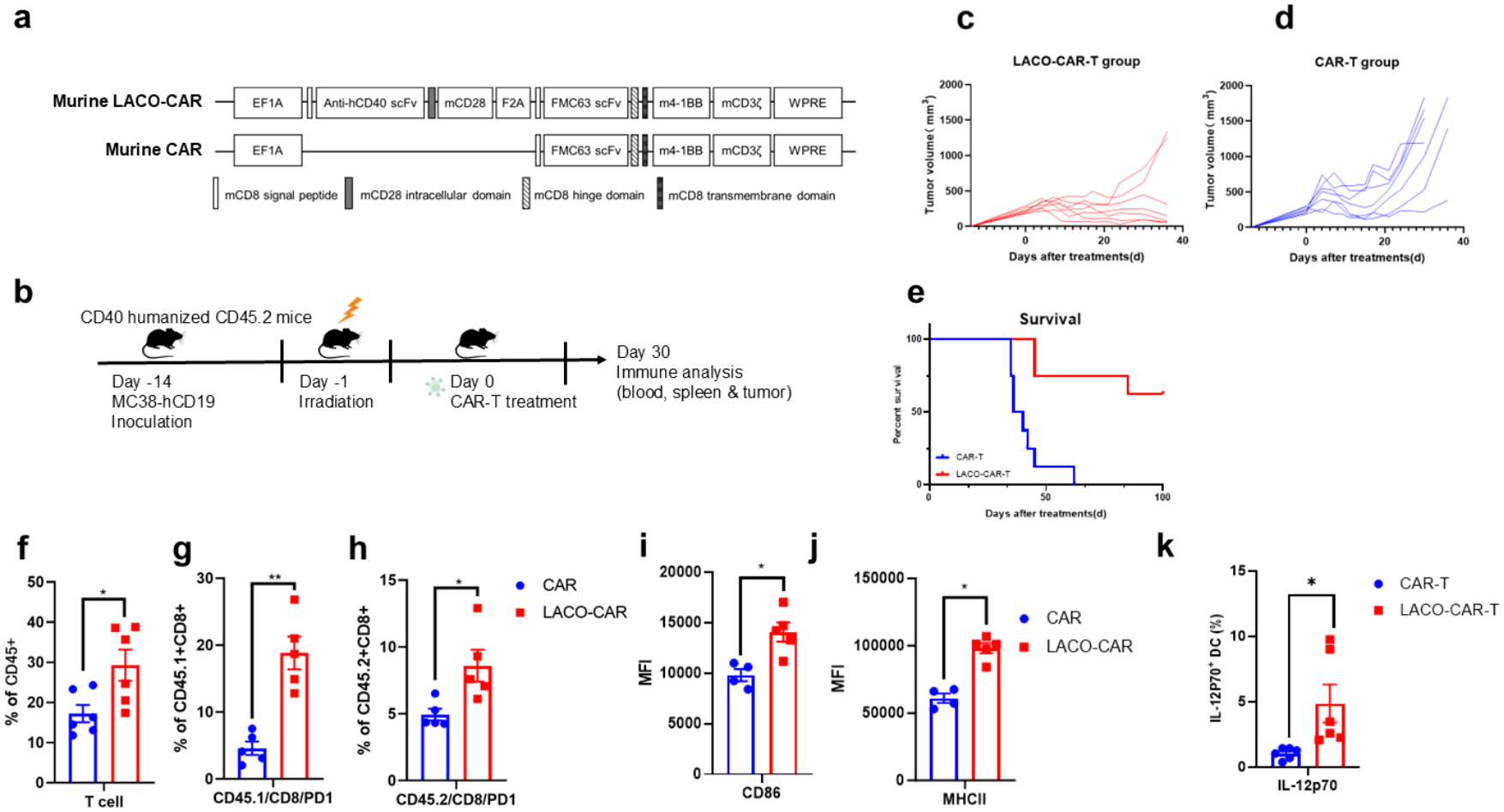
Recipient T cells contribute to tumor control in LACO-CAR-T-treated immunocompetent mice. a, Schematic of the murine LACO-CAR and CAR constructs; FMC63 scFv, the hCD19-targeted single-chain variable fragment. b, Schematic of the in vivo experiments. Humanized CD40/CD45.2 C57BL/6 mice were inoculated with MC38-hCD19 cells and lymphodepleted (4.75 Gy), then received CAR-T or LACO-CAR-T cells (3 × 10⁶, intravenous) prepared from CD45.1 C57BL/6 mice. One cohort was sacrificed on day 30 for flow-cytometry analysis of immune phenotypes in spleen, tumor and blood. c, d, Tumor growth curves for the LACO-CAR-T (c) and CAR-T (d) groups measured by calipers until day 36 after treatment. e, Survival monitored for 100 days in the CAR-T and LACO-CAR-T groups. f, Flow-cytometry analysis of activated T cells (CD3^+^) in tumors after CAR-T or LACO-CAR-T treatment. g, h, PD-1 expression on exogenous (CD45.1^+^) CD8^+^ T cells (g) and endogenous (CD45.2^+^) CD8^+^ T cells (h) in tumors. i, j, Flow-cytometry analysis of CD86 and MHC class II expression (MFI) on tumoral DCs (CD45.2^+^ CD3⁻ B220⁻ CD11c^+^ MHC-II^+^ population). k, Quantification of intracellular IL-12 (IL-12p70) in tumoral DCs. A total of 2 × 10⁶ digested tumor cells were fixed, permeabilized and washed according to the intracellular staining kit (BioLegend), stained with 5 μl PE-IL-12p70 antibody (BD Biosciences) for 30 min on ice, followed by surface staining; IL-12p70 staining was measured in the CD45.2^+^ CD3⁻ B220⁻ CD11c^+^ MHC-II^+^ population.

### LACO-Stim promotes epitope spreading and enriches neoantigen-specific endogenous T cells

To determine whether LACO-CAR-T-induced tumor-infiltrating CD8^+^ T cells recognize antigens beyond the CAR target, we isolated CD8^+^ T cells from tumors on day 28 after treatment, co-cultured them with hCD19-negative parental MC38 cells, and performed an IFN-γ ELISpot assay. IFN-γ-secreting cells were significantly increased in the LACO-CAR-T group compared with the CAR-T group (P < 0.0001), whereas CAR-T treatment did not increase IFN-γ-secreting CD8^+^ T cells relative to untreated controls (P = 0.088) (Fig. 3a, b), providing evidence that LACO-CAR-T treatment enhanced CD8^+^ T-cell responses against endogenous tumor antigens independently of the CAR target. Intracellular cytokine staining after co-culture revealed significantly elevated frequencies of IFN-γ^+^ TNF-α^+^ cells among recipient (CD45.2^+^) CD8^+^ T cells in the LACO-CAR-T group (Fig. 3c, d), consistent with epitope spreading mediated by LACO-Stim-driven APC cross-presentation. Among infused exogenous (CD45.1^+^) CD8^+^ T cells, IFN-γ^+^ TNF-α^+^ frequencies were also increased in LACO-CAR-T-treated mice (Fig. 3e, f), indicating that LACO-Stim augmented T-cell priming through APC reprogramming in both donor and recipient compartments.

**Fig. 3.**
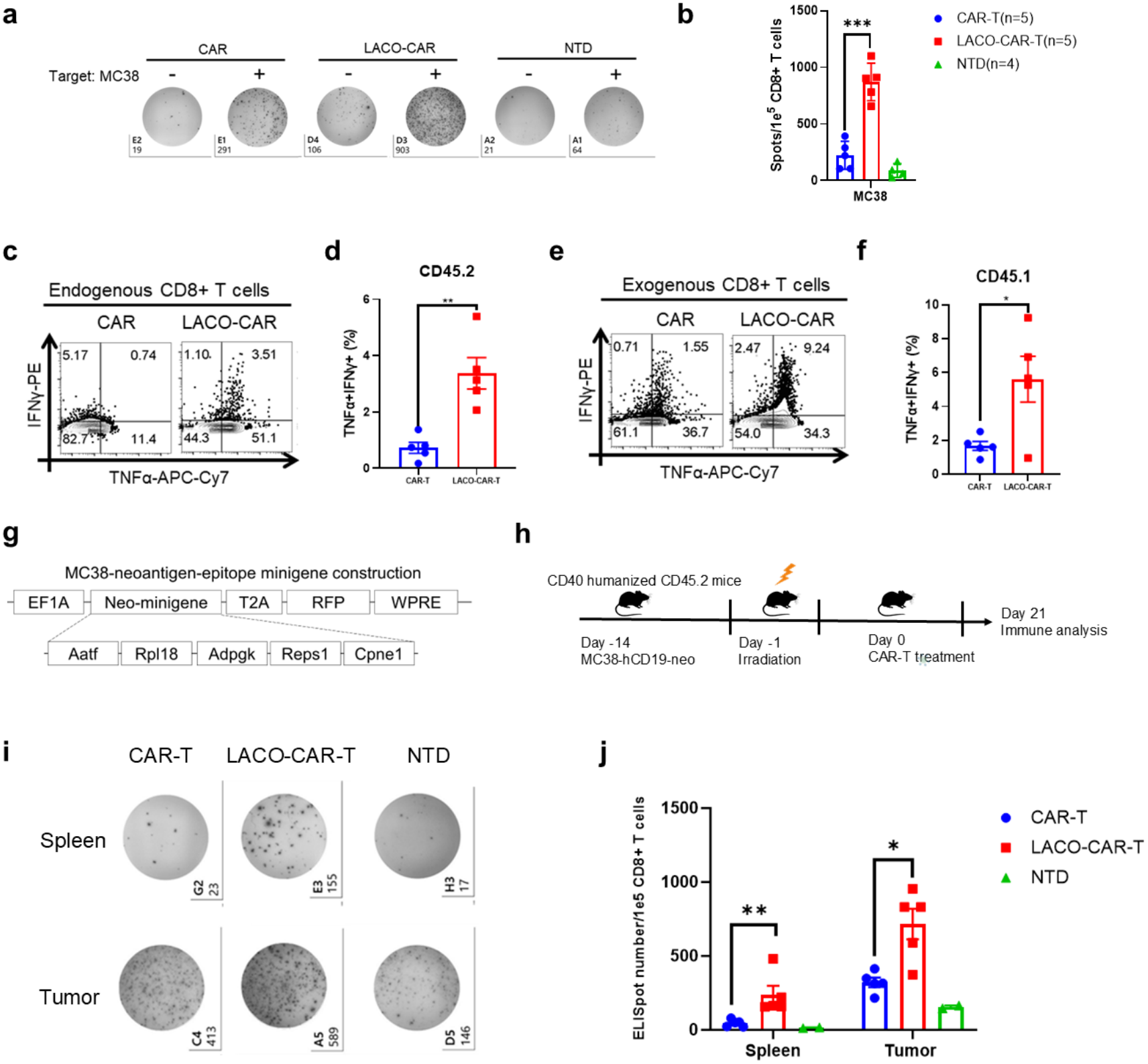
LACO-Stim drives epitope spreading and induces endogenous neoantigen-specific T-cell responses. a, b, Representative IFN-γ ELISpot images of 1 × 10⁵ CD8^+^ T cells sorted from tumors and co-cultured with 1 × 10⁴ MC38 cells. c, d, Flow-cytometry analysis of intracellular TNF-α and IFN-γ expression in recipient CD8^+^ T cells from tumors co-cultured with MC38 for 24 h. e, f, Flow-cytometry analysis of intracellular TNF-α and IFN-γ expression in exogenous CD8^+^ T cells from tumors co-cultured with MC38 for 24 h. g, Schematic of the MC38 neoantigen-epitope minigene:nucleotide sequences encoding each core mutated epitope (Aatf, Rpl18, Adpgk, Reps1 and Cpne1) were connected without spacer, followed by RFP, constructed into a pELNs lentiviral packaging system to transduce MC38-hCD19 tumor lines. h, Schematic of the MC38-hCD19-neo tumor model. Humanized CD40/CD45.2 C57BL/6 mice were inoculated with MC38-hCD19-neo cells (2 × 10⁶, subcutaneous), lymphodepleted (4.75 Gy) and given CAR-T or LACO-CAR-T cells (3 × 10⁶, intravenous) from CD45.1 C57BL/6 mice. Mice were sacrificed on day 21 for immune cell analysis. i, j, Representative IFN-γ ELISpot images and statistics; 1 × 10⁴ mouse bone marrow-derived DCs were pulsed with 10 μg ml⁻¹ of the peptide mixture (Aatf, Rpl18, Adpgk, Reps1 and Cpne1) and co-cultured with 1 × 10⁵ sorted CD8^+^ T cells from spleen or tumor for 24 h.

To identify the epitopes recognized by the induced T cells, we selected five validated neoantigens (Aatf, Rpl18, Adpgk, Reps1 and Cpne1) previously reported in the MC38 model [18]. Because these neoantigens were absent from the MC38 line used in this study [19], we synthesized the epitopes as a minigene construct upstream of red fluorescent protein (RFP) and transduced it into MC38-hCD19 cells to generate MC38-hCD19-neo (Fig. 3g). Low-RFP-expressing lines were sorted and maintained to minimize immunogenicity during tumor establishment. Immune cells from the blood that were co-cultured with neoantigen peptides showed significant immune response toward each peptide after MC38-hCD19-neo vaccination at day 14, indicating that vaccination of hCD40 mice with MC38-hCD19-neo cells elicited neoantigen-reactive T cells (Extended Data Fig. 3a, b). MC38-hCD19-neo tumor-bearing mice were irradiated on day 13 after tumor inoculation and received adoptive transfer of CAR-T or LACO-CAR-T cells (Fig. 3h). Consistent with the parental MC38-hCD19 model, LACO-CAR-T treatment significantly suppressed MC38-hCD19-neo tumor growth relative to CAR-T (Extended Data Fig. 3c, d). In parallel, tumors and spleens were collected on day 21 after treatment. Sorted CD8^+^ T cells were co-cultured with bone marrow-derived murine DCs pulsed with neoantigen peptide mixtures; ELISpot analysis showed significantly higher numbers of IFN-γ-secreting CD8^+^ T cells in the LACO-CAR-T group in both spleen and tumor (Fig. 3i, j). These results confirm that LACO-Stim elicits epitope spreading toward neoantigens, promoting coordinated tumor control by infused CAR-T cells and LACO-Stim-induced T cells directed against antigens beyond the CAR-T target.

To further validate APC-dependent epitope spreading, we tested LACO-Stim-expressing OT-I T cells (LACO-OT-I) against B16-F10-OVA (B16-OVA) tumors in hCD40 mice (Fig. 4a). LACO-OT-I treatment achieved superior tumor control (5 of 7 mice) compared with OT-I monotherapy (2 of 6) (Fig. 4b–d). In a separate experiment, spleen, lymph node and tumor were harvested on day 30 (Extended Data Fig. 4a). Increased Trp2-reactive CD8^+^ T cells, detected by H-2Kb/Trp2 tetramer staining, were found in the spleen, lymph node and tumor of the LACO-OT-I group compared with the OT-I and non-transduced control groups (Fig. 4e, f). CD8^+^ T cells from LACO-OT-I-treated mice also showed enhanced IFN-γ responses against OVA-negative B16-F10 cells ex vivo (Extended Data Fig. 4b–d), confirming epitope spreading to tumor antigens beyond the therapeutic target. Within the tumor myeloid compartment, macrophages and DCs increased while myeloid-derived suppressor cells (MDSCs) decreased (Extended Data Fig. 4e), and activation and maturation markers were upregulated on tumor macrophages and DCs of the LACO-OT-I group (Extended Data Fig. 4f, g).

**Fig. 4.**
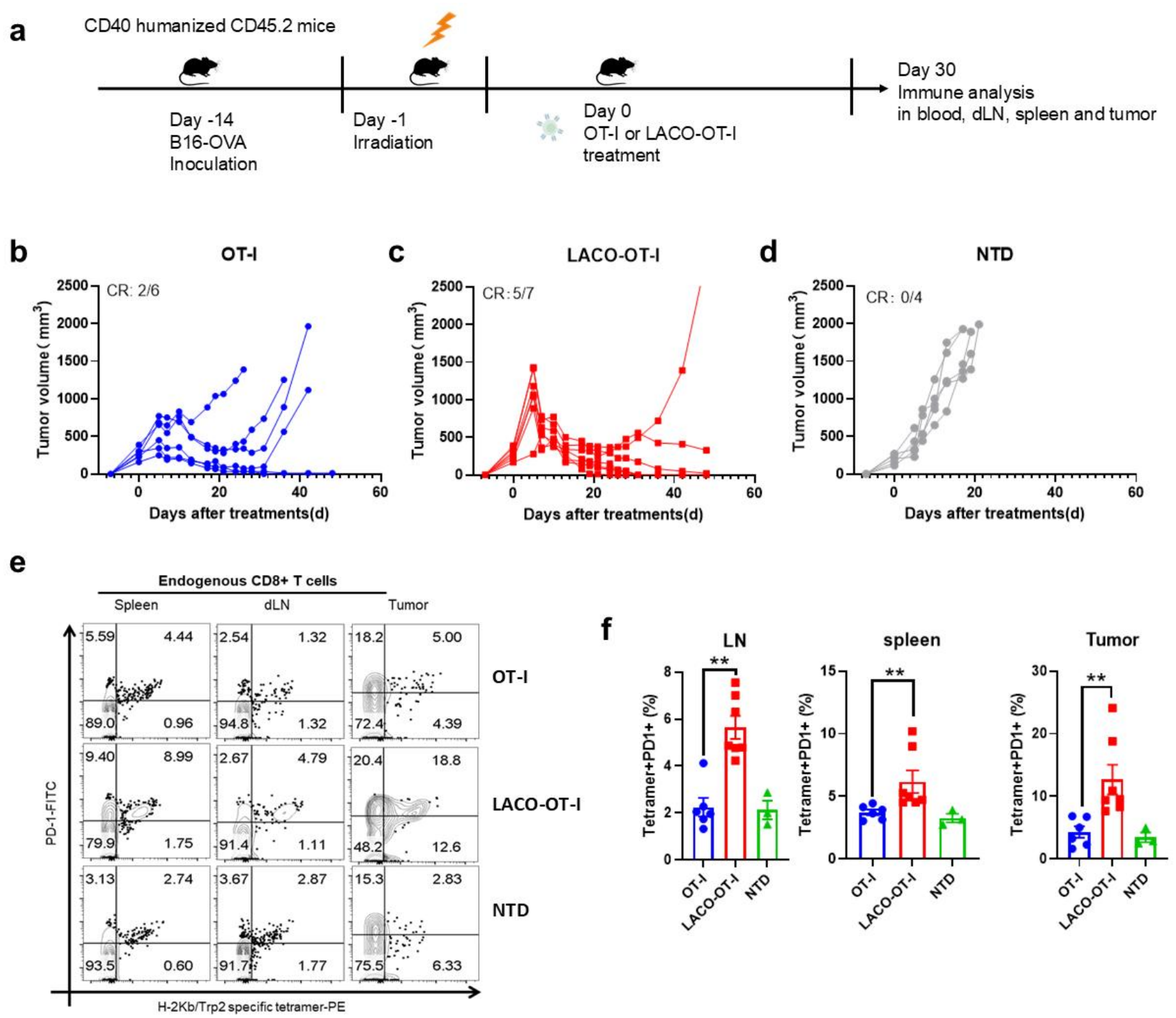
LACO-Stim enhances OT-I T-cell therapy and induces epitope spreading in a B16-OVA melanoma model. a, Schematic of the B16-OVA tumor model. A total of 5 × 10⁵ B16-OVA cells were engrafted into humanized CD40/CD45.2 mice, followed by OT-I or LACO-Stim-expressing OT-I (LACO-OT-I) cells prepared from OT-I transgenic mice; mice were euthanized on day 30 and draining lymph nodes, spleen and tumor were digested for further analysis. b-d, Tumor volume in the OT-I (b), LACO-OT-I (c) and no-treatment (d) groups. e, f, Representative H-2Kb/Trp2 tetramer and PD-1 staining of CD8^+^ T cells from spleen, draining lymph nodes and tumors.

Collectively, these results demonstrate that LACO-Stim remodels the tumor microenvironment and reprograms tumor myeloid compartments, leading to enhanced myeloid-dependent anti-tumor immunity, and that it can serve as a broadly applicable augmentation module for engineered T-cell therapies.

### LACO-Stim reshapes the tumor immune microenvironment at single-cell resolution

To decode how LACO-Stim reshapes host anti-tumor immunity beyond cell-intrinsic co-stimulation, we profiled tumor-infiltrating immune cells from untreated (N), CAR-T-treated (C) and LACO-CAR-T-treated (LC) MC38-hCD19-neo tumors by single-cell RNA sequencing (scRNA-seq; 37,095 cells after quality control, of which 34,163 were CD45^+^ immune cells). Unbiased clustering and reference-based annotation (SingleR/ImmGen) revealed a marked redistribution of the immune landscape (Fig. 5a): T-lineage cells expanded from 11.7% (N) and 13.5% (C) to 39.5% of all profiled cells in the LC group, with CD8^+^ T cells increasing approximately fourfold (6.1% (N), 7.1% (C) to 25.4% (LC)), whereas the macrophage compartment contracted correspondingly (61.3% (N), 61.8% (C) to 18.0% (LC)), yielding an approximately twelvefold increase in the T-cell-to-macrophage ratio (Fig. 5b and Supplementary Table 1).

**Fig. 5.**
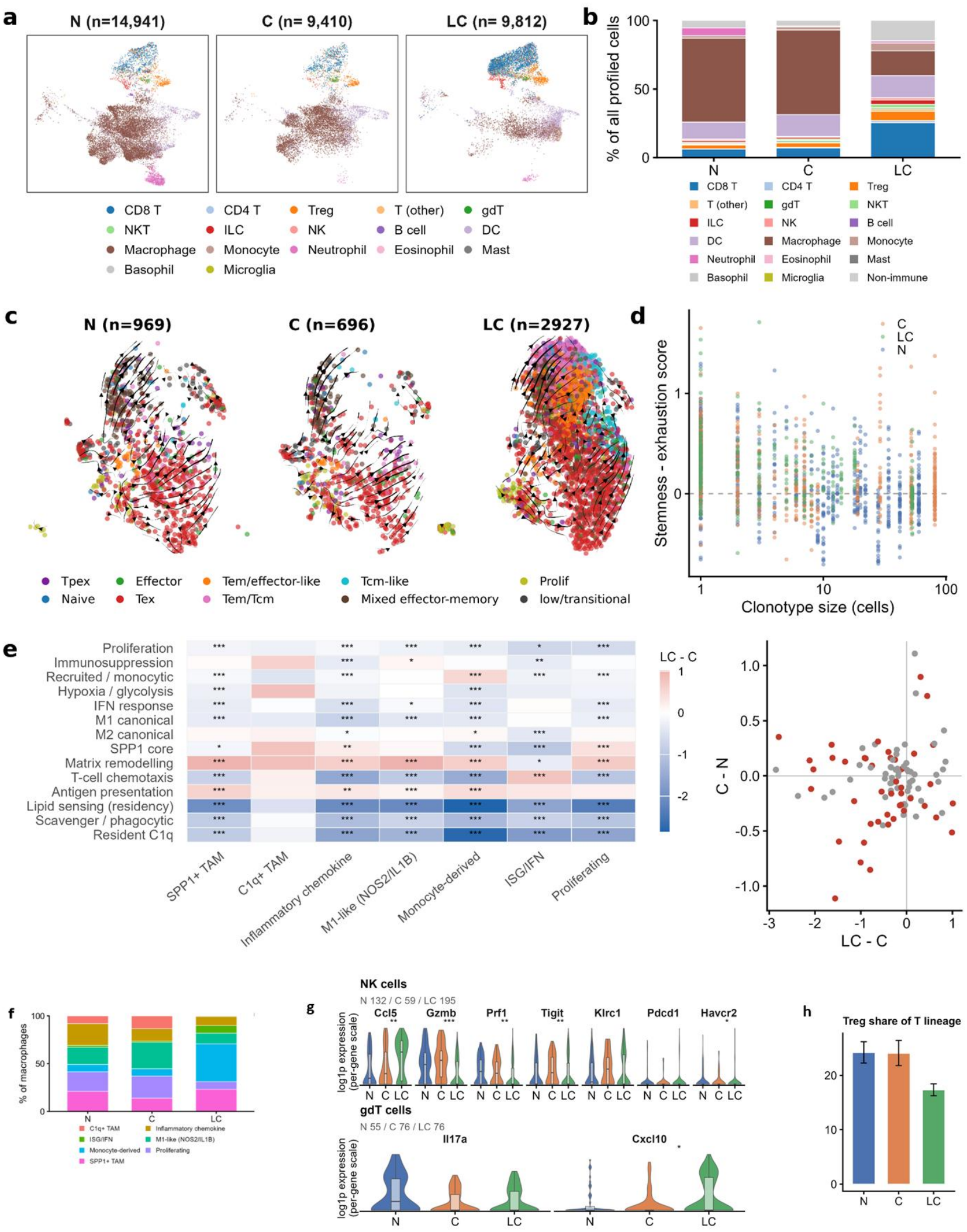
Single-cell transcriptomic and RNA-velocity analysis reveals comprehensive immune remodeling by LACO-CAR-T. a, UMAP visualization of 34,163 CD45^+^ tumor-infiltrating immune cells from non-treatment (N), CAR-T (C) and LACO-CAR-T (LC) groups, colored by cell type (SingleR/ImmGen annotation). b, Cell-type composition across groups (percentages of all profiled cells; n = 37,095; canonical marker-based annotation, with non-immune stromal, endothelial and epithelial cells collapsed into a single grey category). c, RNA velocity of CD8^+^ T cells across treatment groups. UMAP embedding of 4,592 CD8^+^ T cells (N, n = 969; C, n = 696; LC, n = 2,927), with the same coordinates used in all three panels. Cells are colored by differentiation subtype: Naive, Tpex (pre-exhausted), Effector and Tex (terminally exhausted) were assigned from the state annotation, while cells not captured by these states (n = 2,143) were resolved into the finer classes Tem/effector-like, Tem/Tcm, Tcm-like, Mixed effector-memory, Prolif and low/transitional by reference-based decomposition. Streamlines summarize the local RNA velocity field estimated with scVelo (deterministic mode) from spliced/unspliced count ratios and projected onto the UMAP; arrowheads indicate the predicted direction of state transition, and line length and density are a smoothed visual summary of the underlying per-cell velocity vectors rather than a quantitative rate. d, Clonotype size versus the stemness-minus-exhaustion axis score for TCR-annotated CD8^+^ T cells (n = 1,508 cells); dots are single cells colored by group; Spearman r = −0.48, P = 4.8 × 10⁻⁸⁷. e, Reprogramming of the macrophage compartment. Left, per-subtype module-score shifts (log1p-CPM) between LACO-CAR-T and CAR-T alone (heat map; asterisks denote Benjamini-Hochberg-adjusted P values, two-sided Mann-Whitney U test; n values denote cells per subtype, LACO-CAR-T / CAR-T; C1q^+^ TAM, n = 19 in LACO-CAR-T, was excluded from statistics and is greyed out). Right, directional discordance between the two CAR-T contrasts: each point is one module-subtype pair, plotted as the shift between LACO-CAR-T and CAR-T alone (x axis) against the shift between CAR-T and non-treatment (y axis); red points denote pairs significant in both contrasts (both q < 0.05), and 44 of 84 pairs shifted in opposite directions (labelled pairs: resident monocyte-derived, inflammatory and proliferating, and lipid M1-like and SPP1+). Per-module directional consistency is provided in Supplementary Table 5, and the gene lists of all modules are provided in Supplementary Table 10. f, Macrophage subtype composition within the macrophage compartment. g, Expression of cytotoxic-effector, chemokine and exhaustion genes in NK cells (Ccl5, Gzmb, Prf1, Tigit, Klrc1, Pdcd1 and Havcr2) and γδ T cells (Il17a and Cxcl10) across treatment groups (violin plots; per-gene log1p-normalized expression; two-sided Wilcoxon test with Benjamini-Hochberg correction; *q < 0.05, **q < 0.01, ***q < 0.001). h, Treg fraction of T-lineage cells (Wilson 95% CI; chi-square test). RNA-velocity functional-state remodeling of γδ T cells (redirected toward a CXCL10-secreting phenotype), DCs (less inflammatory), Tregs (weakened suppressive function; CTLA-4/CD39/IL-10/PD-1 down). Source data supporting panels a–h are provided in Supplementary Tables 1, 2, 4, 5, 8 and 10.

RNA velocity analysis identified CD8^+^ T cells at different states of differentiation, from naive, Tcf7^+^ precursors of exhausted T cells (Tpex) to terminally exhausted (Tex). Further analysis revealed that LC treatment preserved a larger early-activated pool while reducing terminal exhaustion (Tex 38.0% versus 54.6% in C and 52.9% in N), shifting the stemness-minus-exhaustion axis from −0.209 (N) and −0.155 (C) to −0.067 (LC) (Fig. 5c, d, and Extended Data Fig. 5a; Supplementary Table 2). Mapping CD8^+^ T cells onto a Monocle3 pseudotime axis rooted at the most naive Tcf7^+^ cells showed that LC redirects differentiation away from terminal exhaustion, evidenced by the findings that LC cells were markedly enriched at the early, stem-like end of the trajectory (42.2% versus 21.3% in N and 10.9% in C) and correspondingly depleted at the late/exhausted end (LC 12.7% versus 38.1% in N and 47.9% in C) (Extended Data Fig. 5b; Supplementary Table 3).

Within the myeloid compartment, the dominant change was a broad reprogramming of macrophage states rather than a simple M1/M2 switch (Fig. 5e, f): resident C1q^+^ macrophages collapsed from 13.4% (C) to 0.6% of macrophages in LC, proliferating myeloid cells decreased from 23.1% (C) to 8.1% (LC), and monocyte-derived macrophages expanded from 7.7% (C) to 39.5% (LC), together with an increase in ISG/IFN-responsive states (from 1.3% (C) to 7.9% (LC); Supplementary Table 4). Pan-myeloid module scoring showed concordant erosion of pro-tumoral residency and immunosuppression programs across the myeloid compartment (Fig. 5e, left; directional discordance in Fig. 5e, right), consistent with CD40-agonistic licensing of APCs in situ. Comparison of the two treatment contrasts showed that 44 of 84 module-score pairs shifted in opposite directions between CAR-T alone and LACO-CAR-T (Fig. 5e, right; Supplementary Table 5), with several resident (monocyte-derived, inflammatory and proliferating) and lipid-associated (M1-like and SPP1+) macrophage subtypes among the diverging trajectories, indicating that LACO-Stim CAR-T treatment redirects rather than merely amplifies CAR-T-driven myeloid remodeling. Dendritic cells (DCs) increased among all profiled cells (12.2% to 16.1%; Supplementary Table 1). RNA-velocity indicated a shift toward a less inflammatory DC state in the LC group, with reduced expression of Il1b, Tnf and Cxcl10 (Extended Data Fig. 5c and Supplementary Table 6). RNA-velocity analysis of the monocyte/macrophage compartment confirmed a coordinated shift of this pool toward a less suppressive, less inflammatory and more antigen-presenting state in the LC group: Apoe and IL-10 declined together with Il1b, Tnf and Cxcl10, whereas MHC-II (H2-Aa) expression and the monocyte-derived marker Ly6c2 increased in the LC group (Supplementary Table 6). CellRank absorption probabilities for monocyte-derived macrophages (2,242 cells, all three groups pooled) were highest for the M1-like state (40.3% of total absorption probability), exceeding residency (C1q^+^ TAM, 2.3%), ISG/IFN (0.1%) and SPP1+ TAM (14.9%) programs, with the remaining probability distributed across the monocyte (31.0%), inflammatory-chemokine (7.1%), proliferating (3.3%) and unlabeled (0.9%) states (Supplementary Table 9).

Notably, innate effector populations were transcriptionally reprogrammed (Fig. 5g): NK cells in the LC group downregulated the cytotoxic effectors Gzmb and Prf1 (q = 7.2 × 10⁻⁶ and 2.3 × 10⁻³ versus CAR-T alone) and the inhibitory receptors Tigit and Havcr2 (q = 2.3 × 10⁻³ and 4.7 × 10⁻²), while upregulating the chemokine Ccl5 (q = 3.8 × 10⁻³), with Klrc1 and Pdcd1 unchanged; γδ T cells instead shifted toward a CXCL10-high state (q = 2.1 × 10⁻² versus CAR-T alone), dominated by a bifunctional IL-17^+^ CXCL10^+^ γδT17 subcluster that may act as a chemokine bridge recruiting CXCR3^+^ effector cells rather than undergoing a conventional cytotoxic switch (Fig. 5g and Extended Data Fig. 5d). Regulatory T cells (Tregs) decreased from 24.0% to 17.3% of T-lineage cells (Fig. 5h; Supplementary Table 1), and RNA-velocity analysis further showed that their suppressive function was weakened (CTLA-4, CD39, IL-10 and PD-1 down, Ccr7 up) (Supplementary Table 6).

Cell-cell communication analysis showed that CAR-T alone (C) was the most communication-active arm, carrying the highest MHC-I/MHC-II, PD-L1, Galectin and ADGRE signaling, whereas LACO-CAR-T (LC) reset the network toward the untreated baseline. In the composition-matched analysis, total interaction strength fell from 53.2 in C to 35.1 in LC, comparable to untreated tumors (34.0), whereas in the original (unadjusted) analysis the number of significant ligand-receptor interactions fell from 2,252 to 1,190 (versus 1,497 untreated; Supplementary Table 7), without globally suppressing communication. MHC-I signaling returned to the untreated level (LC 10.8 versus N 9.4; CAR-T peak 19.1) even as per-cell antigen-presentation genes rose above untreated tumors in LC myeloid cells (H2-Ab1 +0.70, H2-Eb1 +0.58 and Cd74 +0.33 log1p-CPM; all q < 1 × 10⁻²⁹), and only the suppressive arm fell below baseline (lower Galectin-9–TIM-3 signaling, with T-cell Pdcd1 (−0.33) and Lgals9 (−0.11) reduced below untreated levels (q = 6.9 × 10⁻¹⁶ and 1.8 × 10⁻⁹), whereas Cd274 was lower than in CAR-T alone (−0.06; q = 5.8 × 10⁻⁶) but not significantly below untreated levels (−0.02; q = 0.12) (Extended Data Fig. 5e and Supplementary Table 7). SPP1- and MIF-centered ligand-receptor pairs dominated the residual LC interactions (Extended Data Fig. 5f). Because CellChat interaction strengths scale with cell number, we repeated this analysis with equalized cell-type composition; the reset toward baseline and the selective Galectin-9–TIM-3 reduction were preserved, whereas the apparent PD-L1 and MHC-II network changes were composition-sensitive and are not interpreted (Extended Data Fig. 5g).

Because LACO-Stim is designed to act through CD40 on antigen-presenting cells, we next asked whether the dendritic-cell (DC) compartment acquired enhanced antigen cross-presentation. Depth-matched module scoring (count-thinned to a common per-cell depth) of the pooled DC compartment (cDC1, cDC2-like and mature DCs) showed that LACO-CAR-T did not expand cross-presentation capacity relative to CAR-T alone (cross-presentation-core module, Δ[LC−C] = −0.01, n.s.; MHC-II presentation, Δ = −0.23, P < 1 × 10⁻²⁶) and mildly reduced cDC1 identity (Δ = −0.22, P = 4 × 10⁻¹³; Batf3, Irf8 and Xcr1 down, H2-Aa −0.24), indicating that LACO-Stim does not act by enlarging the cDC1 subset. Instead, the DC pool shifted away from a tolerogenic, chronically inflamed state: the negative-regulator module fell (NegReg_Tolerogenic −0.07 versus CAR-T; P = 8 × 10⁻⁵; Supplementary Table 6b), together with lower Il1b, Tnf, Ccl5, Apoe, Irf4 and Cd83 (all P < 0.05 versus CAR-T) and lower Il6 than untreated tumors (P = 7 × 10⁻⁹). In parallel, MHC-I peptide-loading machinery rose above non-treatment (Tap1 +0.35, P = 1.4 × 10⁻⁸; Psmb9 +0.29, P = 9.8 × 10⁻⁴; B2m +0.24 versus CAR-T, P = 1.8 × 10⁻⁶) and the IFN/NF-κB transcriptional program was elevated (Stat1 +0.38, P = 6 × 10⁻⁷; Nfkb1 +0.67, P = 1 × 10⁻¹³; Cxcl9 +0.23, P = 3 × 10⁻³ versus CAR-T), without a corresponding increase in basal TLR/cGAS-STING sensor transcripts. Notably, this less-inflammatory DC state was accompanied by marked upregulation of the checkpoint ligands PD-L1 and PD-L2 (Cd274 +0.52 and Pdcd1lg2 +0.48 versus CAR-T, P < 1 × 10⁻¹⁰), an adaptive counter-regulation consistent with preserved antigen-presentation signaling but requiring combination with PD-1 blockade. Thus, LACO-Stim remodels tumor DCs toward a less tolerogenic, MHC-I/IFN-high state rather than by expanding the cross-presenting cDC1 compartment (Extended Data Fig. 5h, i and Supplementary Table 6c).

### LACO-Stim synergizes with PD-1 blockade to prevent immune escape and achieve durable tumor control

Having shown that LACO-Stim recruits both exogenous (CD45.1^+^) and endogenous (CD45.2^+^) T cells into the tumor and licenses them to recognize neoantigens and tumor-associated antigens, we generated a mosaic tumor model by mixing MC38 with MC38-hCD19 cells at a 1:1 ratio and inoculating the mixture into hCD40/CD45.2 mice (Extended Data Fig. 6a). CAR-T monotherapy transiently slowed tumor growth until day 20 after treatment, after which all tumors grew rapidly, compared with the non-transduced (NTD) group. In contrast, most LACO-CAR-T-treated mice achieved sustained tumor control, with tumor growth still well controlled in treated mice (7 of 10), translating to significantly improved progression-free and overall survival (Extended Data Fig. 6b–f). To test whether checkpoint inhibition could further improve treatment, anti-PD-1 antibody was administered from day 7 after CAR-T infusion (twice weekly for 2 weeks) (Fig. 6a). The combination achieved complete tumor control in 5 of 6 mice for LACO-CAR-T plus anti-PD-1, compared with 3 of 6 with LACO-CAR-T alone, whereas CAR-T plus anti-PD-1 and NTD plus anti-PD-1 each achieved complete tumor control in only 1 of 6 mice (Fig. 6b–f). This synergy was corroborated by significantly superior survival in the LACO-CAR-T plus anti-PD-1 combination group (Fig. 6g).

**Fig. 6.**
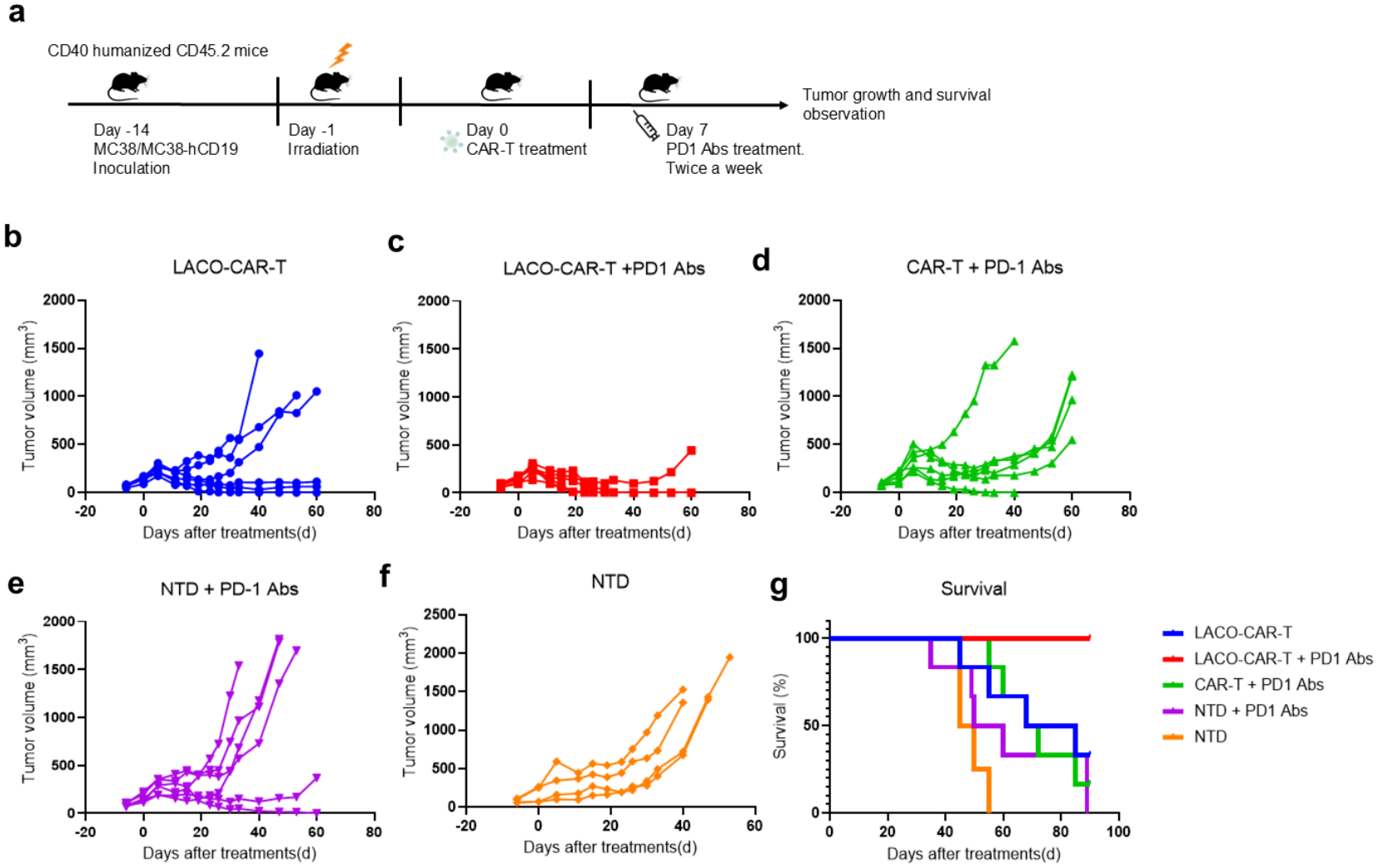
LACO-CAR-T combined with anti-PD-1 blockade achieves durable tumor control in a mosaic tumor model. a, Schematic of the mosaic MC38/MC38-hCD19 tumor model with PD-1 antibody. A total of 5 × 10⁵ MC38 cells were mixed with 5 × 10⁵ MC38-hCD19 cells in 100 μl PBS and engrafted into humanized CD40/CD45.2 mice, followed by irradiation and T-cell treatment. Mice then received intraperitoneal injections of 100 μg PD-1 antibody (BioXCell, BE0146). Tumor growth and survival were monitored. b-f, Tumor volume in the LACO-CAR-T (b), LACO-CAR-T plus anti-PD-1 (c), CAR-T plus anti-PD-1 (d), NTD plus anti-PD-1 (e) and NTD (f) groups. g, Survival statistics of mice in each treatment group.

## Discussion

In our previous work, we introduced a PD-1-CD28 switch receptor into CAR-T cells and found that it potentiates CAR-T expansion and eradicates solid tumors in vivo [20]. PD-L1 is, however, low or absent in most clinical solid tumors, weakening the switch that drives T-cell proliferation [21, 22], and downregulation of the CAR target can aggravate immunosuppression and favor immune escape [23]. To address both limitations, we replaced the PD-1 ectodomain of the switch receptor with an anti-CD40 scFv, retaining the CD28 intracellular domain. CD40 is expressed mainly on APCs and on most solid tumors [24, 25]; both CD40L modification and CD40 agonist antibodies enhance adoptive T-cell therapy through DC activation and the elicitation of endogenous anti-tumor responses [17, 26, 27], yet they leave TME-imposed T-cell exhaustion unresolved. Because CD40L requires trimerization and supramolecular clustering for efficient CD40 engagement, a requirement poorly compatible with the transient, spatially constrained T-cell-APC contacts of solid-tumor stroma [28], we selected a functionally validated agonistic anti-CD40 scFv that, when tethered to the T-cell surface, does not require ligand-like trimerization for CD40 engagement and may therefore provide more consistent APC activation across diverse tumor architectures, while its CD28 domain delivers co-stimulation in cis to the same engineered cell and may avoid the cytokine-release liability reported for membrane-bound CD40L [29]. Licensing APCs alone, however, does not explain how exhausted tumor-infiltrating T cells regain effector function.

Tumor-infiltrating CD8^+^ T cells depend on CD28 co-stimulation more than on any other co-receptor. Exhausted T cells retain TCR signaling but lose the second signal, and PD-1 suppresses them chiefly by recruiting SHP2 to dephosphorylate CD28 rather than the TCR [16]; consistently, rescue of exhausted CD8^+^ T cells by PD-1 blockade is cell-intrinsically CD28-dependent, and the CD8^+^ cells that proliferate in patients on PD-1 therapy are predominantly CD28^+^ [30]. Because B7 ligands are frequently downregulated on tumor cells and limiting on tumor myeloid cells, this node becomes rate-limiting, and supplying it locally is sufficient to drive tumor rejection and to broaden the recognized epitope repertoire [31–33]. Endogenous T cells carry no engineered receptor and must therefore engage CD28 through its natural ligands. A co-stimulatory domain fused within the CAR is not equivalent to ligand-driven CD28 signaling, because its output is shaped by the CD3ζ signalosome and by tonic CAR clustering; native engagement and chimeric switch receptors, which deliver CD28 as a separate, ligand-gated module [20, 34], are therefore the appropriate comparators, which is the configuration adopted by LACO-Stim.

Our in vitro data establish that LACO-CAR-T cells promote DC maturation and M1-like macrophage polarization through CD40 engagement, confirming CD40 agonist activity [35], and these findings were corroborated in vivo by durable tumor control in xenograft and immunocompetent models. Critically, LACO-CAR-T induced epitope spreading, generating endogenous CD8^+^ T-cell responses against neoantigens and other tumor-associated antigens and broadening the anti-tumor spectrum from a single engineered specificity to a polyclonal response [36].

Single-cell transcriptomics and RNA-velocity analysis revealed a coordinated de-inflammation/de-suppression program: LACO-CAR-T drove the monocyte/macrophage pool toward a less suppressive, less inflammatory and more antigen-presenting state (Apoe and IL-10 down, MHC up), rendered DCs less inflammatory, weakened Treg suppressive function (CTLA-4, CD39, IL-10 and PD-1 down) and redirected γδ T cells toward a CXCL10-secreting phenotype. CellChat confirmed that CAR-T alone was the most communication-active arm (highest MHC-I/MHC-II, PD-L1, Galectin and ADGRE signaling), whereas LACO-CAR-T reset the network toward the untreated baseline, preserving antigen-presenting signaling while selectively lowering the suppressive Galectin-9–TIM-3 axis, a microenvironment that is de-inflamed and de-suppressed rather than merely armed. Two caveats apply: CellChat strengths scale with cell number, so key changes were confirmed in a composition-matched analysis (Extended Data Fig. 5g), with PD-L1 network signaling not interpreted because of its composition sensitivity; and with one sample per arm, per-cell tests are pseudo-replicates and are read as supportive. Notably, DCs in LACO-CAR-T tumors also upregulated the checkpoint ligands PD-L1 and PD-L2, an IFN-driven counter-regulation that preserves antigen presentation yet is expected to restrain newly primed CD8^+^ T cells, providing a rationale for the synergy with anti-PD-1 blockade (Fig. 6).

A central question is what sustains the accrual of CD8^+^ T cells and their stem-like, less-exhausted state. Because both infused (CD45.1^+^) and endogenous (CD45.2^+^) T cells increased, this reflects genuine recruitment and priming rather than accumulation of the infused product. The myeloid chemokine arm was, however, not enhanced: T-cell-recruiting ligands on macrophages, monocytes and DCs were unchanged or reduced, and residual chemotactic signal came from stromal and endothelial sources. We therefore propose that the dominant driver is antigen-presentation-coupled co-stimulation within the TME: CD40 agonism licenses tumor macrophages toward an MHC-I/II-high, CD83/NF-κB-low state that presents tumor antigen while extinguishing suppressive cargo (TGF-β1, IL-10, Arg1, SPP1, ApoE and IL-1β), broadening antigen presentation and relieving a critical brake on T-cell function. Within this low-suppression, low-chronic-inflammation niche, tumor-infiltrating CD8^+^ T cells showed the lowest exhaustion burden of the three groups, with a preserved Tcf7^+^Pdcd1⁻ progenitor pool and sustained co-stimulatory receptor availability (CD122, CD28 and 4–1BB). LACO-Stim therefore does not recruit more T cells per se but establishes a permissive niche that licenses and preserves a stem-like, non-terminally exhausted compartment, complementing its cell-intrinsic CD28 co-stimulation. Three caveats should be noted: the analysis rests on a single 28-day time point, so an early expansion that later quiesced cannot be distinguished from a population that never expanded locally; T-cell frequencies are relative compositions from scRNA-seq rather than absolute, marker-defined fractions, concordant in direction but not magnitude with flow cytometry; and DCs did not upregulate MHC-II in the scRNA-seq analysis, despite in vitro maturation, possibly reflecting spatial or temporal constraints on APC licensing that warrant multiplex imaging.

T-cell infiltration increased approximately threefold (Supplementary Table 1) and the repertoire became the most oligoclonal of any group, with 100% group-private top clones, a pattern incompatible with chemokine-gated recruitment. The data instead support direct, CAR-T-intrinsic and antigen-driven T-cell accrual operating in parallel with, but not rate-limited by, macrophage reprogramming. Notably, the SPP1/Arg1 program of SPP1^+^ TAMs was retained even as the IFN-γ-receptor arm declined, suggesting that LACO-Stim remodels macrophage identity without closing the SPP1/Arg1 exit, which may limit durability (Extended Data Fig. 5j).

In addition, LACO-Stim proved complementary to other immunotherapeutic modalities: combination with PD-1 blockade further improved tumor eradication in heterogeneous mosaic tumors, suggesting that relieving T-cell exhaustion amplifies the responses initiated by LACO-mediated APC activation, and in B16-OVA melanoma, LACO-Stim expressed on OT-I TCR-T cells enhanced tumor clearance and induced epitope spreading to Trp2, extending the platform beyond CAR-T.

In summary, LACO-Stim enhances CAR-T-cell function by combining cell-intrinsic co-stimulation with APC-directed microenvironmental reprogramming. This dual-action design may help overcome critical barriers to the treatment of solid tumors and supports further clinical development.

**Extended Data Fig. 1.**
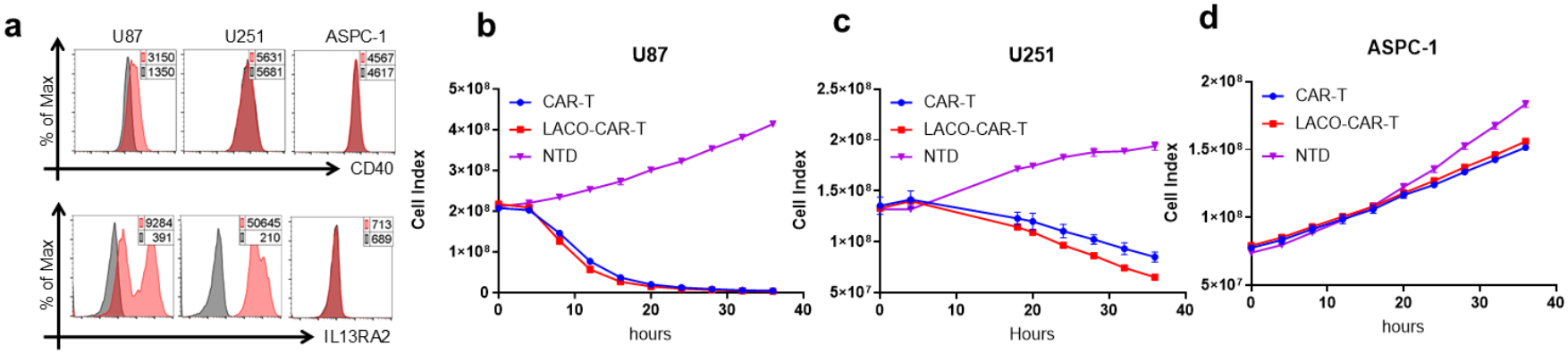
CD40 and IL13Rα2 expression on tumor lines and real-time killing assays. a, CD40 (top) and IL13Rα2 (bottom) expression on U87, U251 and ASPC-1 tumor cell lines. Expression intensity (MFI) is shown in pink numbers compared with isotype staining in grey. b-d, Real-time killing assays of U87 (b), U251 (c) and ASPC-1 (d) tumor lines.

**Extended Data Fig. 2.**
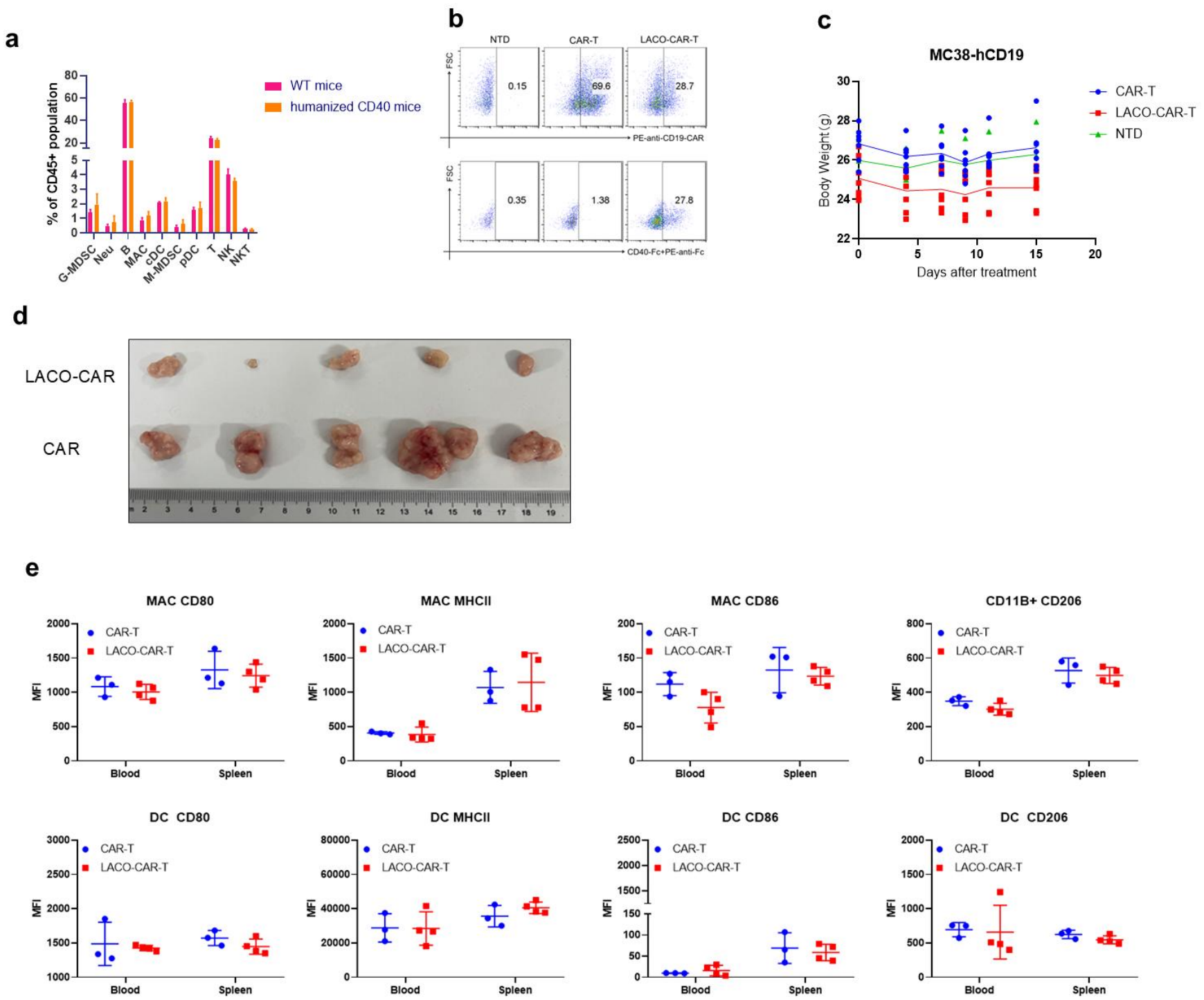
Characterization of humanized CD40 mice and transduced T cells. a, Splenic immune cells, including G-MDSC (CD11B^+^Ly6G^+^Ly6C^low-int^), neutrophil (Neu) (CD11B^+^Ly6G^+^), B cells (CD11B^-^B220^+^), macrophage (MAC) (CD11B^+^F4/80^+^), cDC (CD11C^+^MHCII^+^), M-MDSC (CD11B^+^Ly6G^-^Ly6C^+^), pDC (Ly6C^+^B220^+^), T (CD3^+^NK1.1^-^), NK (CD3^-^NK1.1^+^) and NKT (CD3^+^NK1.1^+^) cells, in wild-type C57BL/6 mice and humanized CD40 C57BL/6 mice. b, CAR and LACO-Stim expression on retrovirally transduced T cells. The CAR positivity rate was corrected against non-transduced T cells, and CAR-positive cell numbers were normalized across groups, so that equivalent numbers of CAR-positive cells were adoptively transferred into mice. c, Body weight of mice after CAR-T, LACO-CAR-T or NTD treatment. d, Tumor photographs at day 30 after CAR-T and LACO-CAR-T treatment. e, CD80, MHC-II, CD86 and CD206 expression on Mφs or DCs from blood or spleen of CAR-T- or LACO-CAR-T-treated mice.

**Extended Data Fig. 3.**
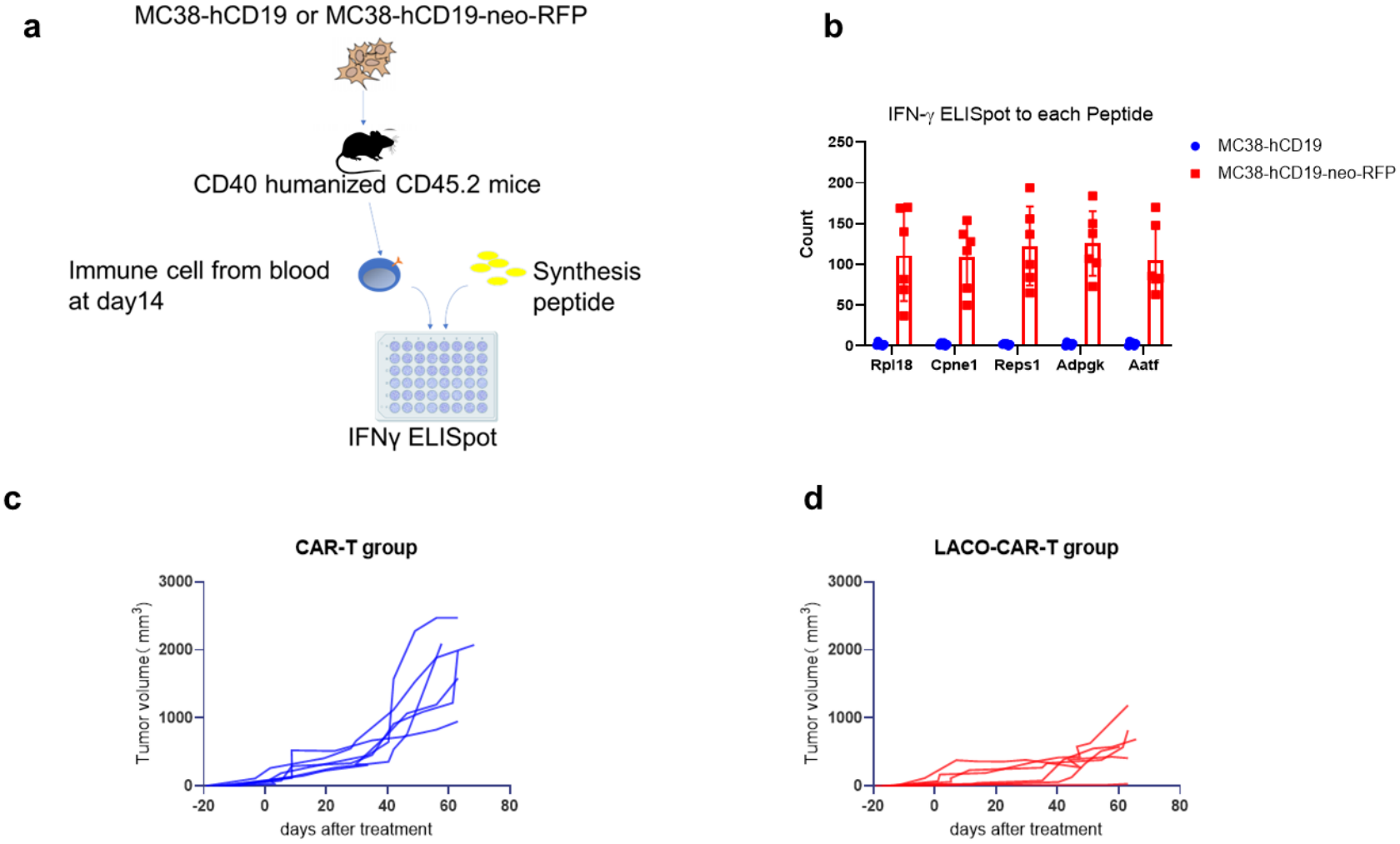
Neoantigen validation and MC38-hCD19-neo tumor model. a, Schematic of immune response toward MC38 neoantigen in tumor model. b, Humanized CD40/CD45.2 C57BL/6 mice were inoculated with MC38-hCD19 or MC38-hCD19-neo cells (2 × 10⁶, subcutaneous), IFN-γ ELISpot assay to detect neo-reactive T cells in blood after tumor engraftment at day 14. A total of 100 μl of mouse blood was collected, red blood cells were lysed, and the cells were co-cultured with synthetic neoantigen peptides (Rpl18, Cpne1, Reps1, Adpgk and Aatf) in an ELISpot plate for 24 h; IFN-γ detection was performed according to the manufacturer’s instructions (Dakewei; Cat#: 2210005). c, d, Tumor growth curves for the CAR-T and LACO-CAR-T groups measured by calipers until day 63 after treatment.

**Extended Data Fig. 4.**
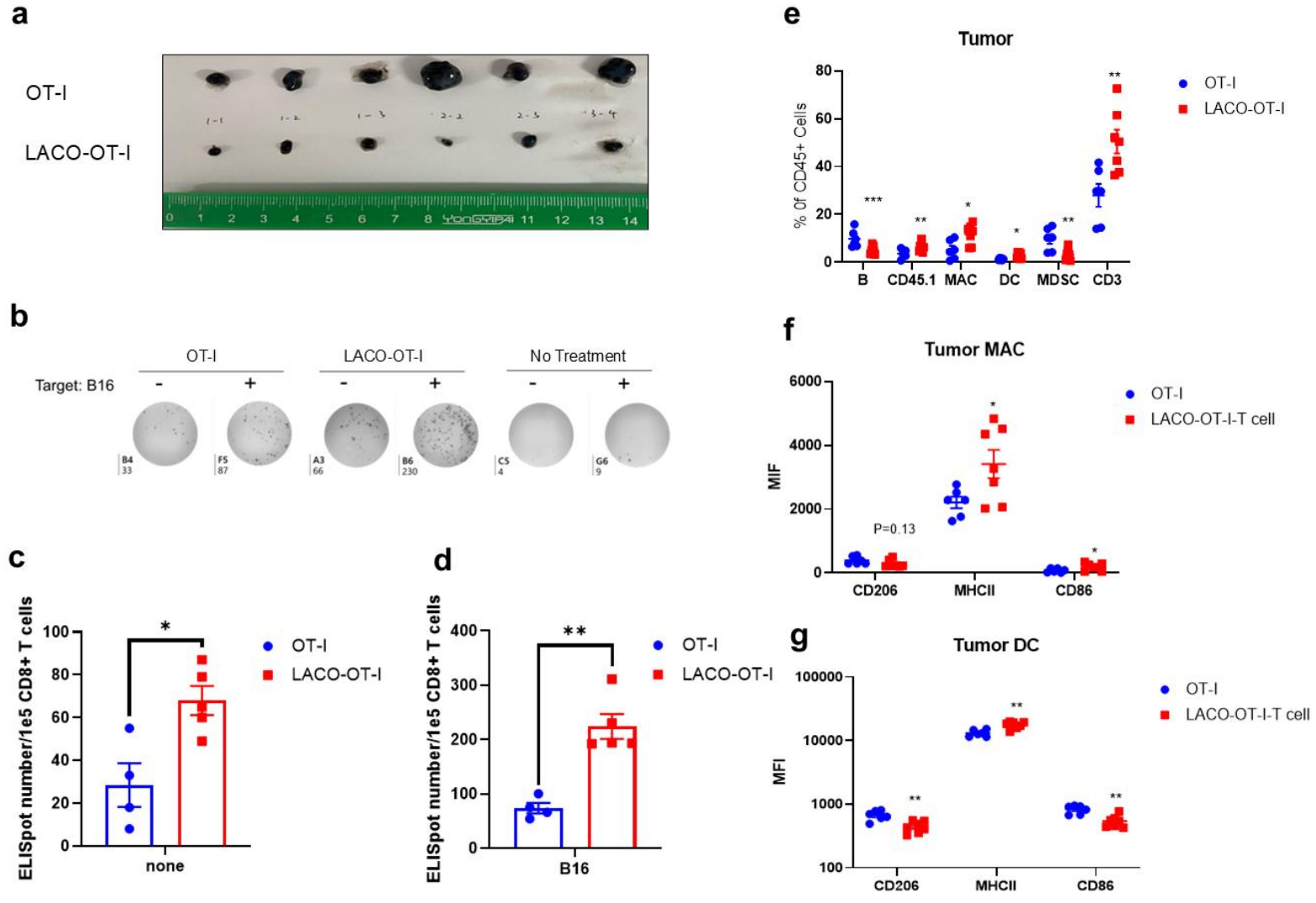
Immune analysis of the B16-OVA model. a, Tumor samples were harvested on day 30. b-d, Representative IFN-γ ELISpot images and statistics of 1 × 10⁵ CD8^+^ T cells sorted from tumors or co-cultured with 1 × 10⁴ parental B16 tumor cells. e, Immune cells in the tumor after treatment. f, g, Activation and maturation markers on tumor Mφs (f) and DCs (g).

**Extended Data Fig. 5.**
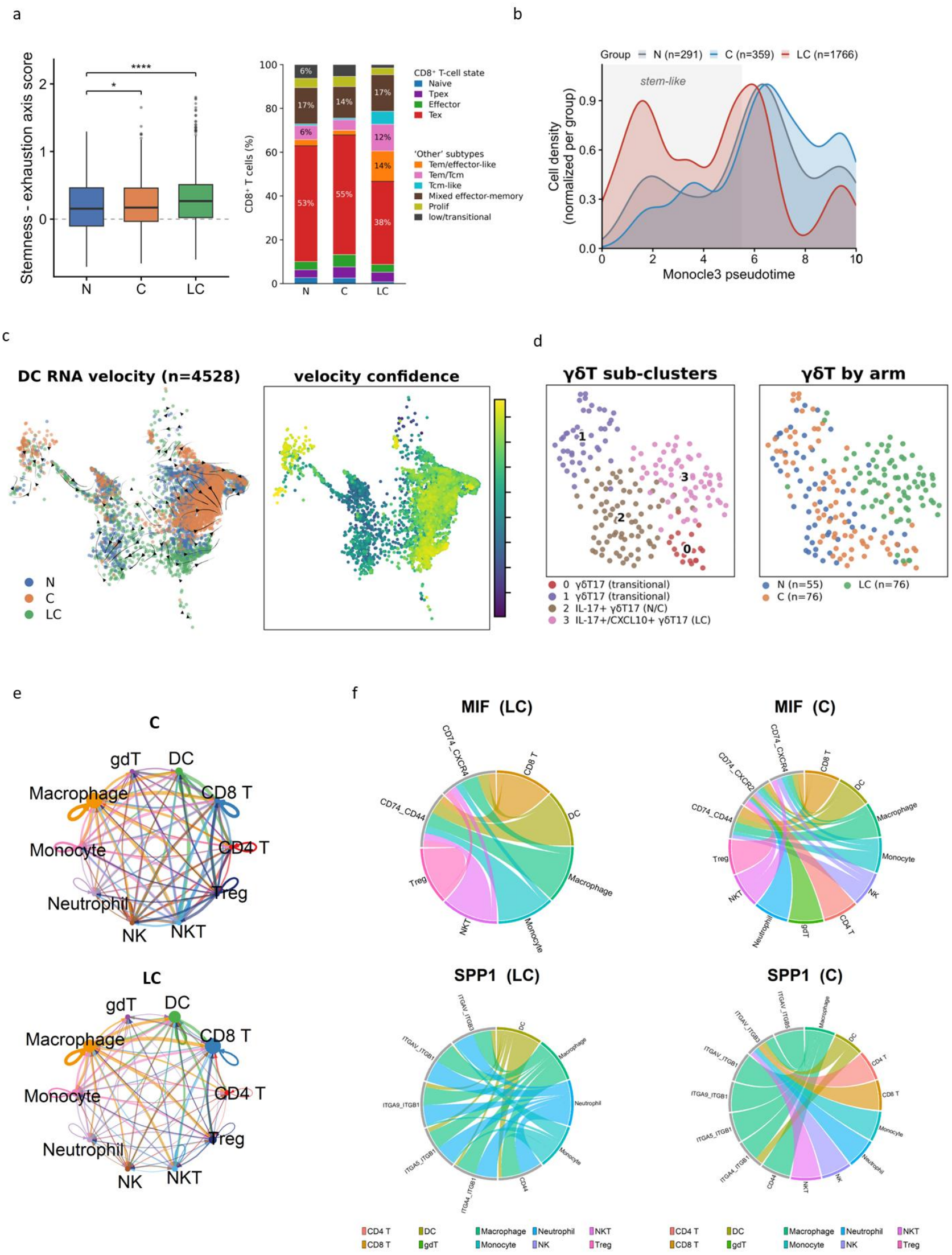
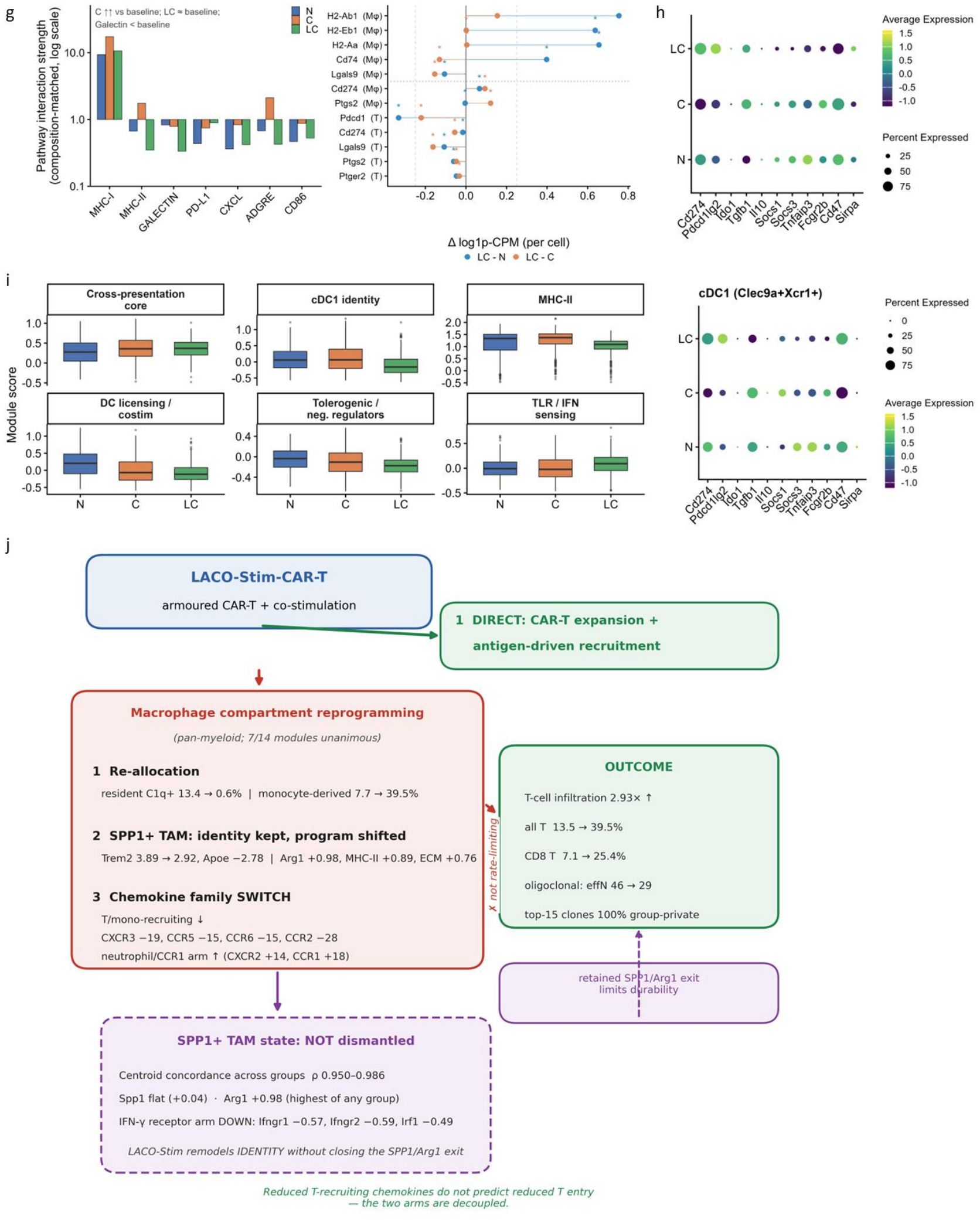
RNA-velocity, trajectory and communication analyses supporting immune remodeling. a, Stemness-minus-exhaustion axis score per group (box plots; two-sided Mann-Whitney U test; *P < 0.05, ****P < 0.0001) (left panel) and 100% stacked bars showing the fraction of CD8^+^ T cells assigned to each state (Naive, Tpex, Effector, Tex) and to the six mechanistic subtypes resolved within the ’other’ pool. Numbers of cells are given below the bars; percentages are shown for segments ≥6%. Colors match the velocity embeddings in **c**. χ² test across the ten categories, P = 2.8 × 10⁻⁷⁸ (right). b, Density of CD8^+^ T cells along the Monocle3 pseudotime axis (rooted at the most naive Tcf7^+^ cells), showing each group’s distribution across the trajectory (normalized within group; N, n = 291; C, n = 359; LC, n = 1,766). LC cells were enriched at the early, stem-like end of the trajectory, whereas control C cells accumulated at the late end. c, DC RNA-velocity streamlines and velocity confidence over the DC UMAP. d, γδ T-cell subclustering and RNA-velocity: the LC group is dominated by a CXCL10-secreting, PD-1-high bifunctional IL-17^+^ γδT17 state (cl3), whereas N/C retain the classic Vδ4 γδT17 (cl2); a Cd8a+ αβ T contaminant (cl0) was excluded. e, CellChat intercellular communication rewiring in C (upper panel) versus LC (lower panel) (circle plots; node size, number of cells; edge width, interaction strength). f, SPP1- and MIF-centered CellChat chord diagrams in the LC and C groups. g, Composition-matched sensitivity analysis of intercellular communication and per-cell statistics: (left) pathway-level interaction strength for key pathways in the composition-matched CellChat analysis relative to the original (proportional-composition) analysis; (right) per-cell log-fold changes (LC versus untreated and LC versus CAR-T) for antigen-presentation and suppressive-axis genes, with Benjamini-Hochberg-adjusted q values. h, Per-gene dot plot of antigen-presentation and negative-regulator/checkpoint genes in the pooled DC compartment (left) and in cDC1 (right), comparing non-treatment (N), CAR-T (C) and LACO-CAR-T (LC). Dot color, mean depth-matched log-normalized expression; dot size, percentage of cells expressing the gene; shaded rows, checkpoint ligands. i, Depth-matched module scores for the pooled DC compartment (cDC1, cDC2-like and mature DC; LC n = 343, C n = 341, N n= 311) across six cross-presentation and licensing modules (CrossPresentation_Core, cDC1_Identity, MHCII_Presentation, DCLicensing_Costim, NegReg_Tolerogenic and TLR_IFN_Sensing); the remaining six modules are reported in Supplementary Table 6b, shown as box plots (two-sided Mann-Whitney U test: *P < 0.05, **P < 0.01, ***P < 0.001, ****P < 0.0001; brackets compare LACO-CAR-T with CAR-T and with non-treatment). Module scores were computed after count-thinning to a common per-cell depth. LACO-CAR-T left the cross-presentation-core module unchanged relative to CAR-T alone (Δ = −0.01, n.s.) and mildly reduced cDC1 identity (Δ = −0.22, P = 4 × 10⁻¹³; Batf3, Irf8 and Xcr1 reduced), but shifted the DC pool toward a less tolerogenic, less inflammatory state (NegReg_Tolerogenic −0.07 versus CAR-T; P = 8 × 10⁻⁵; Supplementary Table 6b) with higher MHC-I peptide-loading (Tap1 +0.35, P = 1.4 × 10⁻⁸ versus non-treatment) and IFN/NF-κB transcriptional activity (Stat1 +0.38 and Nfkb1 +0.67 versus CAR-T, P < 1 × 10⁻⁶). j, Schematic model of LACO-CAR-T-driven T-cell infiltration: an antigen-driven, CAR-T-intrinsic recruitment arm operates in parallel with, but is not rate-limited by, the macrophage chemokine-family switch; the retained SPP1/Arg1 exit of SPP1+ TAMs is proposed to limit durability. Source data for the analyses shown in panels a–i are provided in Supplementary Tables 1–8 and 10.

**Extended Data Fig. 6.**
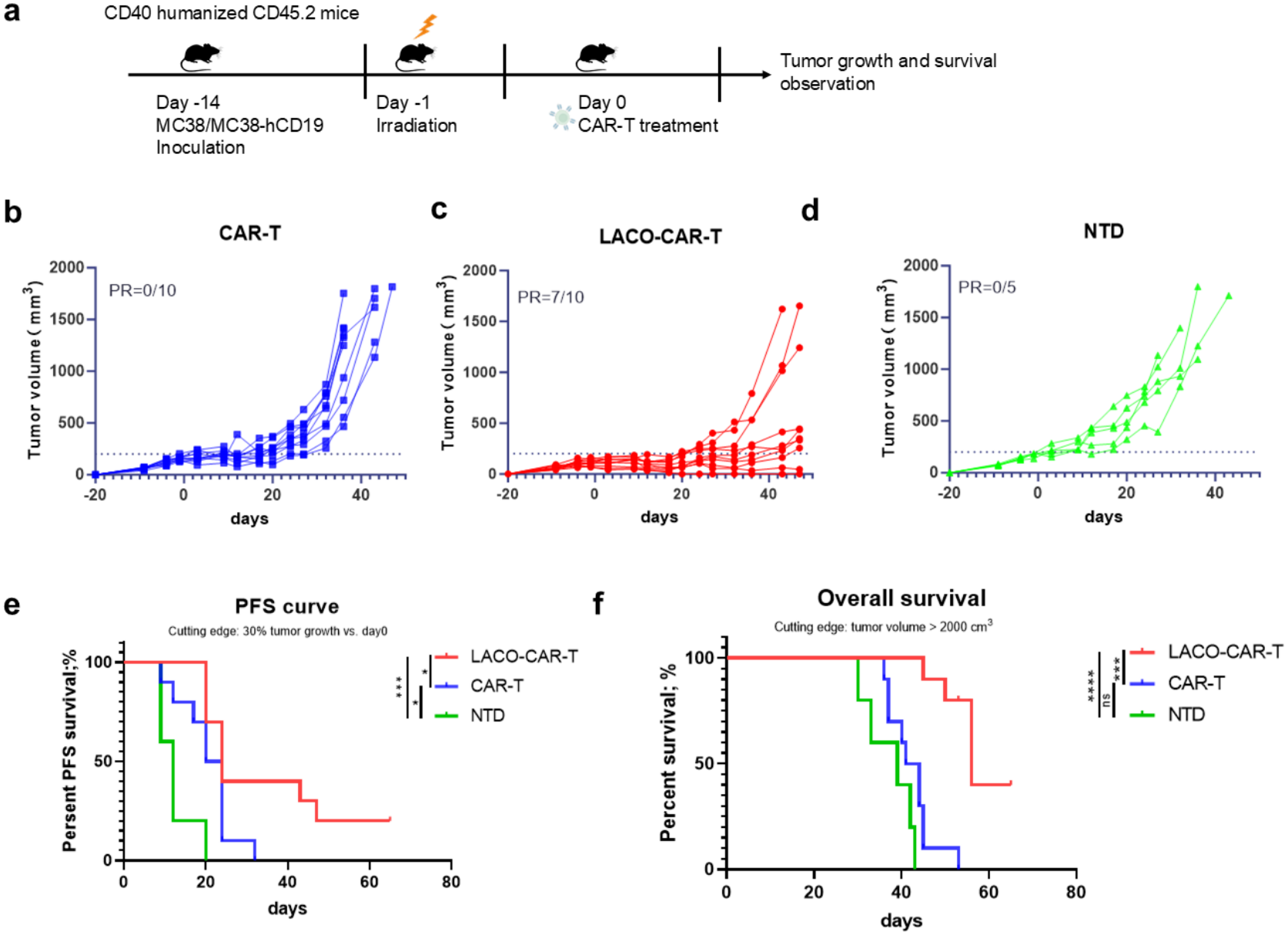
Mosaic MC38/MC38-hCD19 tumor model. a, Schematic of the mosaic MC38/MC38-hCD19 tumor model. Humanized CD40/CD45.2 C57BL/6 mice were inoculated subcutaneously with 5 × 10⁵ MC38 mixed with 5 × 10⁵ MC38-hCD19 followed by lymphodepletion (4.75 Gy) and treatment with 3 × 10⁶ CAR-T or LACO-CAR-T cells intravenously. b-d, Tumor volume in the CAR-T (b), LACO-CAR-T (c) and NTD (d) groups. e, f, Progression-free survival (e) and overall survival (f) of mice in each treatment group.

## Materials and Methods

### Cell lines and cell culture

Human peripheral blood mononuclear cells (PBMCs) were isolated from leukoreduction collars of healthy donors by Ficoll-Paque PLUS density-gradient centrifugation. Single-cell suspensions were obtained from spleen and lymph nodes by mechanical dissociation through 70-μm cell strainers followed by red blood cell lysis with ACK lysis buffer (Gibco). HEK293T (ATCC CRL-3216), U87 MG (ATCC HTB-14), U251 MG (ECACC 09063001), ASPC-1 (ATCC CRL-1682), MC38 (Kerafast ENH204-FP), B16-F10 (ATCC CRL-6475) and B16-OVA cells were maintained in DMEM (Gibco) or RPMI-1640 (Gibco) supplemented with 10% heat-inactivated fetal bovine serum (ExCell Bio or Dongling Bio), 1% GlutaMAX-I (Gibco), 1% non-essential amino acids (Gibco), 1% sodium pyruvate (Gibco), 1% HEPES (Gibco) and 1% penicillin/streptomycin (Gibco) in a humidified incubator at 37 °C with 5% CO2. All cell lines were tested monthly for mycoplasma contamination. Stable reporter cell lines (U87-CBG-GFP, U251-CBG-GFP, ASPC-1-CBG-GFP, MC38-hCD19 and MC38-hCD19-neo-RFP) were generated by lentiviral transduction followed by fluorescence-activated cell sorting on an S3e cell sorter (Bio-Rad).

### Selection and characterization of the anti-CD40 scFv

The anti-human CD40 scFv was obtained by panning a phage display library and screened for binding to recombinant human CD40 by ELISA, followed by a functional screen in which the scFv, expressed as LACO-Stim in CAR-T cells, was tested against DCs for induction of CD80, CD83 and IL-12. The lead clone showed no detectable binding to murine CD40, consistent with the need for hCD40-humanized mice.

### Vector construction and viral production

Lentiviral transfer plasmids were constructed in the pELNs backbone using standard restriction-enzyme-based cloning and Gibson assembly (New England Biolabs). The LACO-Stim element comprised an agonistic anti-human CD40 single-chain variable fragment (scFv) fused to the human CD8α hinge and transmembrane domain (amino acids 138–206) and the human CD28 intracellular domain (amino acids 180–220), linked to the CAR expression cassette via a self-cleaving F2A sequence. The second-generation CAR targeting IL13Rα2 (H08 scFv) contained the human 4–1BB (TNFRSF9) intracellular domain (amino acids 214–255) and human CD3ζ intracellular domain (amino acids 52–163). For the murine system, the anti-human CD19 CAR used the FMC63 scFv with the mouse CD8α signal peptide, hinge and transmembrane, the mouse CD28 proximal intracellular sequence, and the mouse 4–1BB and CD3ζ intracellular domains. The neoantigen minigene construct (neo-T2A-RFP) encoding five dominant MC38 neo-epitopes (Aatf, Rpl18, Adpgk, Reps1 and Cpne1) was synthesized (GenScript) as a single open reading frame.

For lentivirus production, HEK293T cells were seeded at 1.5 × 10⁷ cells per 15-cm dish and transfected 18 h later using PEIpro (Polyplus) in antibiotic-free medium. A mixture of 20 μg transfer plasmid, 7.5 μg pMD2.G, 4.5 μg pRSV-Rev and 6 μg pMDLg-pRRE was incubated with 50 μl PEIpro in 1.5 ml Opti-MEM (Gibco) for 30 min at room temperature before addition to cells. Viral supernatants were harvested at 24 and 48 h, filtered through 0.45-μm PVDF filters (Merck Millipore, SLHV033RS) and concentrated by ultracentrifugation at 25,000g for 2 h at 4 °C in an SW28 rotor (Beckman Coulter). Pellets were resuspended in Opti-MEM, aliquoted and stored at −80 °C.

For retrovirus production, HEK293T cells were transfected with 16 μg of pCL-Eco (NovoPro, V002459) and 24 μg of the MSCV-based retroviral vector using 40 μg PEIpro in antibiotic-free culture medium. Retroviral supernatants were collected at 48 and 72 h post-transfection, filtered, aliquoted, and stored at −80 °C.

### Human CAR-T-cell generation

CD4^+^ and CD8^+^ T cells were isolated from human PBMCs by negative selection using EasySep Human CD4^+^ and CD8^+^ T Cell Enrichment Kit (StemCell Technologies, 19052 and 19053), achieving more than 95% purity as confirmed by flow cytometry. Purified T cells were cultured in complete RPMI-1640 medium containing 100 IU ml⁻¹ recombinant human IL-2 (R&D Systems, 202-IL). T cells were activated with CTS anti-CD3/CD28 Dynabeads (Gibco, 40203D) at a 3:1 bead-to-cell ratio for 24 h, followed by lentiviral transduction at a multiplicity of infection (MOI) of 3:1. Dynabeads were removed 72 h after transduction using a DynaMag magnet (Invitrogen). T cells were maintained at 1 × 10⁶ cells per ml and expanded for 10–15 days until the average cell volume stabilized at approximately 400 μm³ as measured by a Coulter counter (Beckman). Expanded CAR-T cells were used immediately or cryopreserved in serum-free freezing medium (CST, 07930).

### Mouse CAR-T and OT-I T-cell generation

Mouse CD3^+^ T cells were purified from spleens of CD45.1^+^ C57BL/6 mice using the EasySep Mouse T Cell Isolation Kit (StemCell Technologies, 19851). Cells were cultured in complete RPMI-1640 medium supplemented with 50 IU ml⁻¹ recombinant mouse IL-2 (R&D Systems, 402-ML) and 50 μM 2-mercaptoethanol (Sigma, M3148). T cells were activated on 24-well plates pre-coated with 2 μg ml⁻¹ anti-mouse CD3 (eBioscience, 16–0031-85) and 2 μg ml⁻¹ anti-mouse CD28 (eBioscience, 16–0281-85) antibodies for 24 h. Activated T cells were transduced by spinoculation with retroviral supernatant at 1,000g for 90 min at 32 °C in the presence of 10 μg ml⁻¹ polybrene (Sigma, H9268). Cells were cultured for 4–5 days before use. OT-I T cells were generated using the identical protocol with OT-I TCR transgenic CD45.1^+^ donor mice.

### Monocyte-derived dendritic cell and macrophage differentiation

Human CD14^+^ monocytes were isolated from PBMCs using the EasySep Human CD14 Positive Selection Kit II (StemCell Technologies, 17858). For monocyte-derived dendritic cell differentiation, purified monocytes were cultured in complete RPMI-1640 medium supplemented with 100 ng ml⁻¹ recombinant human GM-CSF (R&D Systems, 215-GM) and 20 ng ml⁻¹ recombinant human IL-4 (R&D Systems, 204-IL) for 9 days, with fresh cytokines added every 2–3 days. For macrophage differentiation, monocytes were cultured with 10 ng ml⁻¹ recombinant human GM-CSF for 7 days. Mouse bone marrow-derived dendritic cells were generated by flushing bone marrow from femurs and tibiae of hCD40 transgenic C57BL/6 mice and culturing with 20 ng ml⁻¹ recombinant mouse GM-CSF (R&D Systems, 415-ML) for 7 days. Mouse bone marrow-derived macrophages (BMDMs) were differentiated with 50 ng ml⁻¹ recombinant mouse M-CSF (R&D Systems, 416-ML) for 7 days.

### In vitro functional assays

For cytotoxicity assays, 1 × 10⁵ CAR-T, LACO-CAR-T or non-transduced (NTD) T cells were co-cultured with 1 × 10⁴ CBG-GFP-expressing target cells in 96-well flat-bottom plates (Corning) in triplicate. Target-cell viability was monitored every 4 h for 72 h using an IncuCyte S3 live-cell imaging system (Sartorius). Percent specific lysis was calculated as [1 - (GFP count in experimental well / GFP count in target-alone well)] × 100.

For activation assays, 1 × 10⁵ T cells were co-cultured with 1 × 10⁵ tumor cells in 96-well U-bottom plates. CD107a surface mobilization was assessed after 4 h in the presence of GolgiStop (BD Biosciences, 554724; 1:1,500 dilution) and anti-CD107a antibody (BD Biosciences). Cytokines (IL-2 and IFN-γ) in 24-h supernatants were quantified by ELISA using DuoSet kits (R&D Systems).

For APC activation assays, 1 × 10⁵ human monocyte-derived DCs or Mφs were co-cultured with 1 × 10⁵ T cells for 24 h. Culture supernatants were analyzed by LEGENDplex bead-based immunoassay (BioLegend, 740446) on an Attune NxT flow cytometer (Thermo Fisher). Cells were surface-stained for CD80 (BioLegend, 305220), CD83 (BioLegend, 305310), CD86 (BioLegend, 374206), HLA-DR (BioLegend, 307652), CD206 (BioLegend, 321122) and CD163 (BioLegend, 333606). For phagocytosis assays, Mφs, U87-CBG cells and rested NTD, CAR-T or LACO-CAR-T cells were co-cultured in 96-well flat-bottom plates for 18 h. Mixed cells were stained with anti-human CD45 (BioLegend, 304032) and anti-human CD3 (BioLegend, 317318) antibodies, followed by 7-AAD live/dead reagent (BD Biosciences, 559925). The macrophage population was gated as 7-AAD⁻CD45^+^CD3⁻, and GFP intensity in this population was analyzed using FlowJo software.

### Epitope spreading assays

Tumors were dissociated using the mouse Tumor Dissociation Kit (Miltenyi Biotec, 130-096-730) on a gentleMACS Octo Dissociator. Tumor cells were depleted using PE-conjugated anti-human CD19 antibody (BioLegend, 302208) with the EasySep PE Positive Selection Kit II (StemCell Technologies, 17667). CD8^+^ T cells were enriched using the EasySep Mouse CD8a Positive Selection Kit II (StemCell Technologies, 18953). For IFN-γ ELISpot, 1 × 10⁵ CD8^+^ T cells were co-cultured with 1 × 10⁴ MC38 cells or neoantigen bone marrow-derived DCs in pre-coated ELISpot plates (Dakewei, 2210005) for 24 h. Spots were counted using an ImmunoSpot S6 Analyzer (Cellular Technology Ltd). For intracellular cytokine staining, 1 × 10⁵ CD8^+^ T cells were co-cultured with target cells for 24 h in the presence of GolgiPlug (BD Biosciences, 555029; 1:1,000), then surface-stained for CD45.1 (BioLegend, 110722), CD45.2 (BioLegend, 109822) and CD8 (BioLegend, 100734), followed by fixation/permeabilization and intracellular staining for IFN-γ (BioLegend, 505806) and TNF-α (BioLegend, 506306).

### Tumor models

All mouse studies used 6–8-week-old male and female mice. For human xenograft studies, NOD.Cg-PrkdcscidIl2rgem1Smoc (NM-NSG-001) mice were inoculated subcutaneously with 1 × 10⁶ U87-CBG-GFP cells in 100 μl PBS. After 14 d, mice received intravenous injection of 1 × 10⁷ CAR-T, LACO-CAR-T or NTD cells. Tumor burden was assessed by bioluminescence imaging (IVIS Lumina Series III, PerkinElmer) at 15 min after intraperitoneal injection of 200 μl D-luciferin (Gold Biotechnology; 15 mg ml⁻¹ in PBS).

For immunocompetent studies, humanized CD40 transgenic mice (hCD40/CD45.2, C57BL/6 background) were generated as described in the Results. Mice were inoculated subcutaneously with 1 × 10⁶ MC38-hCD19 or 2 × 10⁶ MC38-hCD19-neo cells. Established tumors (d 10–14, approximately 50–100 mm^3^) were lymphodepleted by total body irradiation (4.75 Gy, 0.8 Gy min⁻¹, RS-2000 X-ray irradiator; Rad Source Technologies). The next day, 3 × 10⁶ CAR+ T cells were injected intravenously via the lateral tail vein in 100 μl PBS. Mosaic tumor models used a mixture of 5 × 10⁵ MC38 and 5 × 10⁵ MC38-hCD19 cells at a 50:50 ratio. For anti-PD-1 combination, 100 μg of anti-mouse PD-1 antibody (BioXCell, BE0146, clone RMP1-14) or isotype control (BioXCell, BE0089) was administered intraperitoneally as indicated. For B16-OVA melanoma models, mice received 5 × 10⁵ B16-OVA cells subcutaneously, irradiation on day 13 and 3 × 10⁶ OT-I or LACO-OT-I cells on day 14 post tumor inoculation.

Tumor size was measured every 3–4 d using digital calipers; tumor volume was calculated as (length × width^2^)/2. Mice were euthanized when body weight loss exceeded 15%, tumor volume exceeded 2,000 mm³, or humane endpoints were met.

### Histology and immunohistochemistry

Tumors were fixed in 10% neutral-buffered formalin (Sigma, HT501128) for 48 h at room temperature, washed with PBS, and stored in 70% ethanol at 4 °C. Tissues were paraffin-embedded, sectioned at 5 μm, deparaffinized and subjected to heat-induced antigen retrieval in citrate buffer (10 mM sodium citrate, 0.05% Tween 20, pH 6.0) at 95 °C for 20 min. Sections were blocked with 10% normal goat serum, incubated with anti-human CD45 antibody (BioLegend, 304002; 1:200) overnight at 4 °C, and detected using HRP-conjugated secondary antibody (Vector Laboratories, MP-7402) and DAB substrate (Vector Laboratories, SK-4105). Sections were counterstained with hematoxylin and scanned at 20× on an Aperio CS2 system (Leica Biosystems). CD45^+^ cell density was quantified using Visiopharm 2023.09 software.

### Flow cytometry

Single-cell suspensions were prepared from tumors, spleen and lymph nodes. Fc receptors were blocked with anti-mouse CD16/32 (BioLegend, 101330; 1:200) or Human TruStain FcX (BioLegend, 422302) for 10 min at 4 °C. Surface staining was performed with fluorochrome-conjugated antibodies at optimal concentrations for 20 min at 4 °C. Intracellular staining was performed after fixation and permeabilization with Cytofix/Cytoperm (BD Biosciences, 554722). Tetramer staining used PE-conjugated H-2K^b^/Trp2 tetramer (ACRO Biosystems) for 30 min at room temperature. Samples were acquired on an Attune NxT (Thermo Fisher Scientific) or LSRFortessa (BD Biosciences) and analyzed with FlowJo v10.8.1 (BD Biosciences) using fluorescence-minus-one (FMO) gating controls.

### Single-cell RNA sequencing

CD45^+^ tumor-infiltrating immune cells were enriched using the EasySep Mouse CD45 Positive Selection Kit (StemCell Technologies, 18967) and processed on a Chromium Controller (10x Genomics) with the Chromium Next GEM Single Cell 3’ Reagent Kit v3.1 (10x Genomics, PN-1000268). Libraries were sequenced on an Illumina NovaSeq 6000 at approximately 50,000 reads per cell and processed with Cell Ranger v7.1.0 (10x Genomics) aligned to mm10. Downstream analysis was performed with scanpy (v1.12) and scVelo (v0.3.4) in Python. After quality control (cells with fewer than 200 or more than 5,000 detected genes, more than 20% mitochondrial reads or more than 10% hemoglobin reads excluded; 37,095 cells retained), samples were integrated with Harmony and clustered using the Leiden algorithm (resolution 0.6, 24 clusters). Cell types were annotated with SingleR using the ImmGen reference (18 classes). RNA velocity was computed with scVelo (deterministic model) from spliced/unspliced counts generated by velocyto and visualized as UMAP streamlines; CellRank was used for fate probabilities and Monocle3 for CD8^+^ T-cell pseudotime analysis rooted at Tcf7^+^ cells. Cell-cell communication was inferred with CellChat. Differentially expressed genes were identified by the Wilcoxon rank-sum test (|log2 fold-change| > 0.25, Benjamini-Hochberg-adjusted P < 0.05) and curated gene-program module scores were computed per cell. All scores, modules and gene panels used in the single-cell analysis, with their complete gene lists, scoring method and provenance, are listed in Supplementary Table 10. Cell-composition percentages were computed on all cells; expression- and module-based statistics were computed after uniform down-sampling to 3,000 UMIs per cell to control for sequencing-depth differences between groups; the depth-matched DC gene-level analysis reported in Supplementary Table 6c used a 4,000-UMI target. Raw and processed data have been deposited in GEO under accession number GSE350560.

### Statistical analysis

Data are presented as mean ± s.d. or s.e.m. as indicated in figure legends. Statistical analyses were performed using GraphPad Prism 9.0 (GraphPad Software). Comparisons between two groups used unpaired two-tailed Student’s t-test after confirmation of normal distribution (Shapiro-Wilk test) and equal variance (F-test). Comparisons among three or more groups used two-way ANOVA. Tumor growth curves were analyzed by two-way repeated-measures ANOVA with Sidak’s correction. Survival was analyzed by the Kaplan-Meier method with the log-rank (Mantel-Cox) test. For scRNA-seq, the Wilcoxon rank-sum test with Benjamini-Hochberg correction was applied. A two-sided P < 0.05 was considered significant. Sample sizes were determined based on prior experience; no statistical power analysis was performed a priori. Randomization used computer-generated sequences; investigators were blinded to group allocation during data collection where feasible.

### Study approval

All animal experiments were approved by the Institutional Animal Care and Use Committee of Shanghai Model Organisms Center (SMOC-IACUC, approval no. 2025-0004; approved 20 January 2025). Mice were housed under SPF conditions on a 12 h light–dark cycle with *ad libitum* access to food and water, and all efforts were made to minimize suffering. Procedures complied with the ARRIVE guidelines and relevant institutional and national animal welfare regulations.

Peripheral blood was collected from healthy adult donors recruited through Milecell (Shanghai) Biotechnology Co., Ltd, with written informed consent (including permission for publication of de-identified data) and ethics approval from the providing institution. PBMCs were isolated and de-identified before downstream generation of dendritic cells, macrophages or CAR-T cells. All procedures were performed in accordance with the Declaration of Helsinki and applicable regulations.

Tumor cell lines were authenticated by STR profiling and tested negative for mycoplasma. All lentiviral and retroviral vector work was performed in a BSL-2 facility under institutional biosafety approval, in accordance with the requirements of the Shanghai Pudong New Area Health Commission (filing certificate no. SYS310115837363).

### Data availability

The single-cell RNA-seq and TCR-seq data generated in this study have been deposited in the Gene Expression Omnibus (GEO) under accession number GSE350560 and will be released upon publication; until then, the data are available from the corresponding author upon reasonable request. All other data supporting the findings of this study are available within the paper and its Supplementary Information or from the corresponding author upon reasonable request. Supplementary Tables 1–10 are provided as a single Excel workbook.

### Code availability

Custom code used for single-cell RNA-seq and TCR-seq analysis is available from the corresponding author upon reasonable request.

### Use of AI

Computational analyses for single-cell RNA-seq and parts of the manuscript text were assisted by large language model tools. All code, analyses, results, and text were verified and approved by the authors, who are solely responsible for the content of this work.

## Acknowledgements

We would like to thank Jintao Feng, Xiangming Feng and Yating Du for technical assistance.

## Funding

Research funding was provided by:

1. National Key Research and Development Program of China (2023YFC3403800)

2. Zhejiang Provincial Innovation and Entrepreneurship Team (2023R02012)

## Author contributions

Writing of the original draft was performed by Z.Z. and Y.Z. Conceptualization of the experiments was performed by Z.Z., F.Y., G.Z., J.W., C.H., X.L. and Y.Z. Review and editing of the manuscript was performed by Z.Z., X.L. and Y.Z.

